# A ten-year community reporting database reveals rising coyote boldness and associated human concern in Edmonton, Canada

**DOI:** 10.1101/2022.10.18.512552

**Authors:** Jonathan J. Farr, Matthew J. Pruden, Robin Glover, Maureen H. Murray, Scott A. Sugden, Howard W. Harshaw, Colleen Cassady St. Clair

**Author notes:** **Corresponding Author** Colleen Cassady St. Clair B 522 Biological Science Building, 11355 - Saskatchewan Drive, University of Alberta, Edmonton, Canada T6G 2E9.

## Abstract

In cities throughout North America, sightings of coyotes (*Canis latrans*) have become common. Reports of human-coyote conflict are also rising, as is the public demand for proactive management to prevent negative human-coyote interactions. Effective and proactive management can be informed by the direct observations of community members, who can report their interactions with coyotes and describe the location, time, and context that led to their interactions. To assess the predictors of human-coyote conflict, we used a web-based reporting system to collect 9,134 community-supplied reports of coyotes in Edmonton, Canada, between January 2012 and December 2021. We used a standardized ordinal ranking system to score each report on two indicators of human-coyote conflict: coyote boldness, based on the reported coyote behaviour, and human perceptions about coyotes, determined from the emotions expressed by reporters. We assigned greater conflict scores to behaviours where coyotes followed, approached, charged or contacted pets or people, and to perceptions where reporters expressed fear, worry, concern, discomfort or alarm. Using ordered logistic regression and chi-square tests, we compared conflict scores for each response variable to spatial, temporal and contextual covariates. Our analysis showed that coyotes were bolder in less developed open areas and during the pup rearing season, but human perceptions were most negative in residential areas and during the dispersal season. Reports that mentioned dogs or cats were more likely to describe bolder coyote behaviour, and those that mentioned pets or children had more negative perceptions about coyotes. Coyote boldness and human perceptions both indicated rising human-coyote conflict in Edmonton over the 10 years of reporting.

## INTRODUCTION

The coyote (*Canis latrans*) is a common example of an urban-adapted species that is also the largest carnivore that is common in cities across North America (Magle et al. 2019; Schell et al. 2020). Coyotes thrive in urban areas largely by avoiding interactions with humans (Mowry et al. 2020, Drake et al. 2021) while benefitting from reduced competition with other predators (Prugh et al. 2009), less human persecution in urban compared to rural areas (Collins and Kays 2011), and abundant urban food resources such as rodents, garbage, compost and fruit trees (Fedriani et al. 2001, Murray et al. 2015a, Sugden et al. 2021). Urban coyotes can potentially also improve human quality of life by regulating populations of rodents and smaller predators like feral cats (Crooks and Soule 1999; Gerht et al. 2013), supporting a sense of connection with nature (Cox and Gaston 2018), and providing an aesthetic enjoyment that is often inherent in seeing wild animals (Soulsbury and White 2015). For these reasons, many people living in cities tolerate, and even appreciate, urban coyotes (Soulsbury and White 2015, Sponarski et al. 2018). However, over the past two decades, there have been increasing reports of bold and aggressive interactions between urban coyotes and people that are indicative of human-coyote conflict and reduce the tolerance of people for populations of urban coyotes (Baker and Timm 2017, Poessel et al. 2017, Draheim et al. 2019). A better understanding of the circumstances associated with conflict could inform approaches to coyote management and public education to support human-coyote coexistence in urban areas.

The level of conflict between humans and coyotes can be assessed in two broad ways: the presence of coyote behaviours, such as boldness or aggression, associated with conflict, and human perception of conflict, indicated by people’s fear or worry about coyotes (Table 1). Coyote behaviours that indicate conflict are often studied retroactively based on the factors associated with attacks, which represent the highest level of human-coyote conflict (White and Gehrt 2009, Baker and Timm 2017). Coyote attacks on pets are usually attributed to predation or the defense of territories or dens (Gehrt et al. 2013, Poessel et al. 2017, Nation and St. Clair 2019). Attacks on humans are rare, but typically generate substantial media attention and degrade public tolerance (Carbyn 1989, Alexander and Quinn 2011, Draheim et al. 2019). These attacks are often preceded by elevated bold or aggressive behaviour, which is frequently caused by increasing coyote habituation to people (Baker and Timm 2017) and associated food conditioning (Lukasik and Alexander 2011). Assessing the factors associated with bold coyote behaviour, which refers to their tendency to approach or interact with people or pets, can help mitigate human-coyote conflict before it occurs at its highest level in the form of attacks.

**Table 1.** Key terms and definitions relating to human-coyote interactions in Edmonton, Alberta.

| **Term** | **Definition** |
| --- | --- |
| Conflict | Occurs when coyotes present a threat, real or perceived, to the well-being of people, pets or property. |
| Coexistence | Co-occurrence of humans and coyotes that minimizes conflict. |
| Sighting | Observation of a coyote at a distance with no interaction. |
| Encounter | Interaction between a coyote and a person/pet at close range. Depending on coyote boldness and human concern, encounters may be reported as positive, if conflict does not occur, or negative, if conflict occurs. |
| Coyote boldness | Ordinal scale indicator of the level of human-coyote conflict determined from the reported coyote behaviour/response towards people or pets. |
| Human concern | Ordinal scale indicator of the level of human-coyote conflict determined from the human perception of coyotes expressed in the report. |

Human perceptions of coyotes may be positive, neutral or negative, with negative perceptions typically relating to concern of harm to themselves, children, or pets when they see or interact with a coyote. Negative perceptions of coyotes, which are associated with conflict, may not align with the actual risk of a coyote attack, but still reduce public tolerance of coyotes, and, consequently, public attitudes towards various forms of wildlife management and policy (Sponarski et al. 2018; Draheim et al. 2019). When attacks on people occur or attacks on pets are prevalent, the public demand for lethal management typically increases with rising negative perceptions of coyotes (Baker and Timm 2017, Draheim et al. 2019). These circumstances make the perception of risk by the public, as indicated by human perceptions about coyotes, an informative metric for coyote management and one for which correlates might also be studied to guide proactive, non-lethal management efforts to minimize potential conflict.

Reports of encounters between humans and coyotes that describe coyote behaviour and human perceptions can both be examined with spatial, temporal and contextual variables via explanatory models. Spatial variables used to predict negative interactions can help identify high-conflict areas, especially when measured at relevant scales to support management actions (Delsink et al. 2013, van Bommel et al. 2020). For example, knowing which areas have low, moderate or high probabilities of interactions between humans and black bears (*Ursus americanus*) allows for management resources to be more efficiently allocated to improve coexistence (Merkle et al. 2011). Temporal variables may apply to scales that range from diel, through seasonal, to inter-annual, again with the goal of predicting when human-wildlife conflict is most likely to occur (Morehouse and Boyce 2017, Soulsbury 2020) so that they can be mitigated more effectively. Additional information for understanding conflict includes contextual variables such as the presence of pets or children, off-leash dogs, and poor health of individual animals (Poessel et al. 2013, Murray et al. 2015; Olson et al. 2015), which influence both coyote behaviour and human perceptions.

The importance of these spatial, temporal and contextual variables for predicting coyote behaviour and human perceptions of conflict can be advanced with large, long-term datasets collected by community members on public reporting interfaces. Such databases allow researchers to gather information over many years and large geographic areas, while simultaneously engaging and educating members of the public (Weckel et al. 2010, Frigerio et al. 2018). Previous studies of this sort have collected voluntary reports of coyote activity using public surveys (Weckel et al. 2010), city reporting databases (Lukasik and Alexander 2011, Poessel et al. 2013), websites (Wine et al. 2015, Mowry et al. 2020), and mobile phone apps (Mueller et al. 2019, Drake et al. 2021). Past analyses of these datasets have shown that reporting varies across levels of urban development, land cover types, and coyote seasons (e.g., breeding, pup-rearing, dispersal; Weckel et al. 2010, Poessel et al. 2013), as well as with human demographic variables such as household income and education (Wine et al. 2015, Mowry et al. 2020). The studies that have assessed different levels of conflict from public reports, based on coyote behaviour, have shown that bold coyote behaviour is most prominent during the coyote pup rearing season (Lukasik and Alexander 2011, Drake et al. 2021). These studies also found more negative interactions in areas where coyotes consume more anthropogenic food (Lukasik and Alexander 2011) and when coyotes were encountered farther from roads (Drake et al. 2021). These studies have provided foundational information that can be applied in cities across North America to facilitate human-coyote coexistence, but, to date, community reports have not yet been used to examine changes in interactions over time. Therefore, extensive and long-term data can provide invaluable information for the additional quantification of the specific spatial, temporal and contextual variables associated with conflict-indicative coyote behaviour and human perceptions about coyotes.

In this study, we used over 9,000 community science reports collected over 10 years from a website in Edmonton, Canada to develop two indicators of human-coyote conflict (Table 1): coyote boldness towards people or pets, based on the reported coyote behaviour, and the human concern about coyotes, determined from the perceptions of coyotes expressed by study participants. Then, we used spatial, temporal and contextual information in the reports to construct exploratory models with the goal of identifying correlates of higher conflict scores for each of coyote boldness and human concern. By revealing the predictors of bold behaviour by coyotes and greater concern about coyotes, we hope to support proactive coyote management and effective public education. In turn, these tools could facilitate sustainable coexistence of humans and coyotes to maximize the ecosystem services, including public appreciation, provided by urban coyote populations throughout North America.

## METHODS

### Study area

This study occurred in Edmonton, Alberta, Canada (53.54728°N, 113.50068°W), which has an area of 684 km^2^ and a population of 976,223 based on the 2019 City of Edmonton Municipal Census (https://www.edmonton.ca/city_government/facts_figures/municipal-census-results). A combination of large area and moderate population density makes Edmonton one of the most sprawling cities in North America, with large areas of undeveloped land (City of Edmonton 2018). Edmonton has warm summers (Jun-Aug daily average = 16.7°C) and cold winters (Dec-Mar daily average = −9.7°C; Environment and Climate Change Canada 2018). The city is bisected by the North Saskatchewan River valley and several large ravines, which form a network of minimally developed natural areas that provide abundant habitat for coyotes and other wildlife (Figure 1).

**Figure 1.**
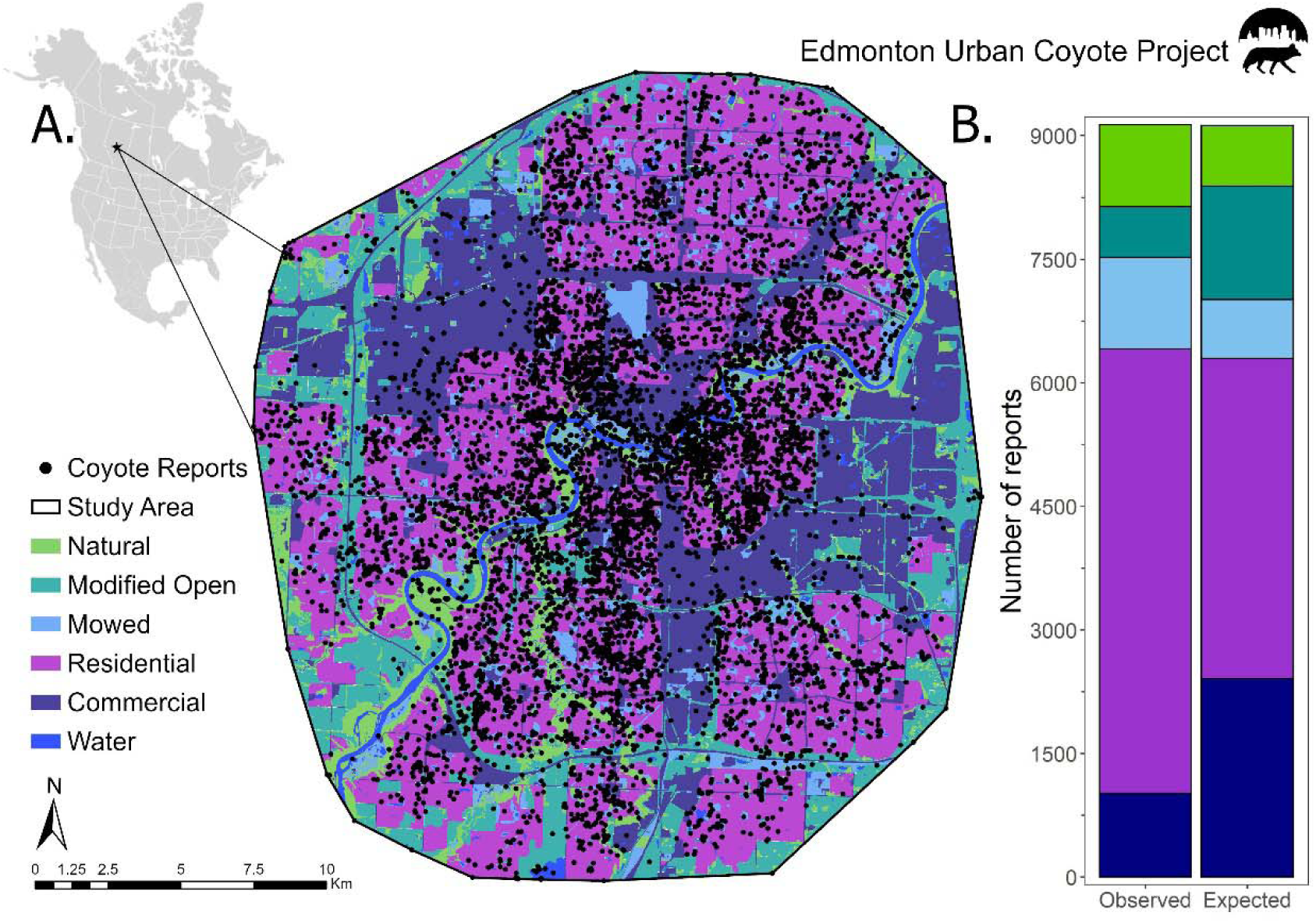
Distribution of coyote reports across Edmonton, Canada (A) and across land cover categories (B). Reports were collected from 2011-2021 through the Edmonton Urban Coyote project website and included the location of the coyote sighting or interaction.

### Report collection

Beginning in September 2010, members of the public were able to voluntarily report coyote sightings or encounters through a web-based platform on the Edmonton Urban Coyote Project website (https://www.edmontonurbancoyotes.ca/reportsighting.php). We promoted the website opportunistically during media interviews, public lectures, and social media posts, as well as through word of mouth, on labels attached to wildlife cameras in the city, and via a link on the City of Edmonton website (https://www.edmonton.ca/residential_neighbourhoods/pets_wildlife/Coyotes.aspx) that was added in 2019.

When submitting a report, participants were asked to provide the location, nearest road intersection of the report location, date, and time of day. The website included a map interface to allow reporters to precisely locate their report by placing a pin on the map. Time of day was submitted by reporters using a drop-down menu with the option to select either hourly times between 5 AM and midnight, or a general time window (dawn, morning, afternoon, evening, or night). Participants were also asked to specify whether their report was a “sighting,” defined as an observation of a coyote at a distance with no interaction, or “encounter,” defined as an interaction with a coyote at close range. Reporters were invited to provide free-form comments, as well as their name and contact information. For the *N* = 3,366 reports that did not include map coordinates, we determined them *post hoc* based on the reported nearest street intersection and other information in the comments (e.g., if a specific park or building was named). To encourage participation, no registration or login was required.

### Extraction of response variables and contextual variables from reports

Most reports (96.8 %, *N* = 8,859) included optional comments with further details about the human-coyote interaction, including information on coyote behaviour, the participant’s perception of coyotes, and various contextual factors. We groomed the database to remove duplicate entries and spam reports, removed the names and contact information of reporters, and then recruited a team of 30 volunteers and undergraduate students who read and classified the comments in each report following an online form and standardized protocol contained in Appendix 1. For each 100 reports that a volunteer or undergraduate classified, one author (JJF) randomly assessed 10 reports to ensure that the protocol was appropriately followed. To further assess the repeatability between report classifiers, JJF randomly selected and blindly re-classified 100 reports. We assessed inter-observer agreement by calculating the percentage of reports that generated the same classifications for each of the seven variables extracted with the classification form.

For reports with comments that described coyote behaviour, volunteers assessed the degree of boldness on a scale from one (ran away) to nine (made physical contact with pets or people). We later simplified these categories into a four-point ordinal scale of coyote boldness defined as avoidance, indifferent, bold and aggressive behaviours (Table 2). We classified human concern about coyotes from reports where participants expressed their perceptions about coyotes to form a three-point scale from the explicit presence of words that conveyed positive (e.g., beautiful), neutral (e.g., curious or not scared), or negative (e.g., scared) emotions or attitudes (Table 3). We also classified report comments for the presence of five contextual variables: (1) the human activity occurring at the time of the report (e.g., walking, cycling, driving), (2) the presence or mention of vulnerable individuals (children, dogs or cats), (3) if dogs present were leashed or off leash, (4) the number of coyotes observed, and (5) any mention of the reporter’s interpretation of coyote health status (e.g., mangy).

**Table 2.** Distribution of coyote reports across coyote boldness ordinal scale values.

| **Coyote Boldness**<br>**(ordinal scale**<br>**value)** | **Description and form classification as**<br>**coyote behaviour** | **Number of**<br>**Reports** | **Percentage of**<br>**Total**<br>**Reports** |
| --- | --- | --- | --- |
| **Avoidance (1)** | Ran away | 645 | 7.2% |
|  | Walked away | 348 | 3.8% |
| **Indifferent (2)** | Did not appear to notice or care about people | 1105 | 12.2% |
|  | Watched the person | 381 | 4.2% |
|  | Vocalized at the person | 46 | 0.5% |
| **Bold (3)** | Followed or stalked pets or people | 464 | 5.1% |
|  | Approached pets or people | 218 | 2.4% |
| **Aggressive (4)** | Chased or charged pets or people | 194 | 2.1% |
|  | Physical contact with pets or people | 139 | 1.5% |
| **Sighting (N/A)** | Reports were submitted as sightings or comments indicated no interaction between people and coyote(s) | 4770 | 52.2% |
| **Unknown (N/A)** | Reports were submitted as encounters, but coyote boldness could not be determined | 824 | 9.0% |

**Table 3.**
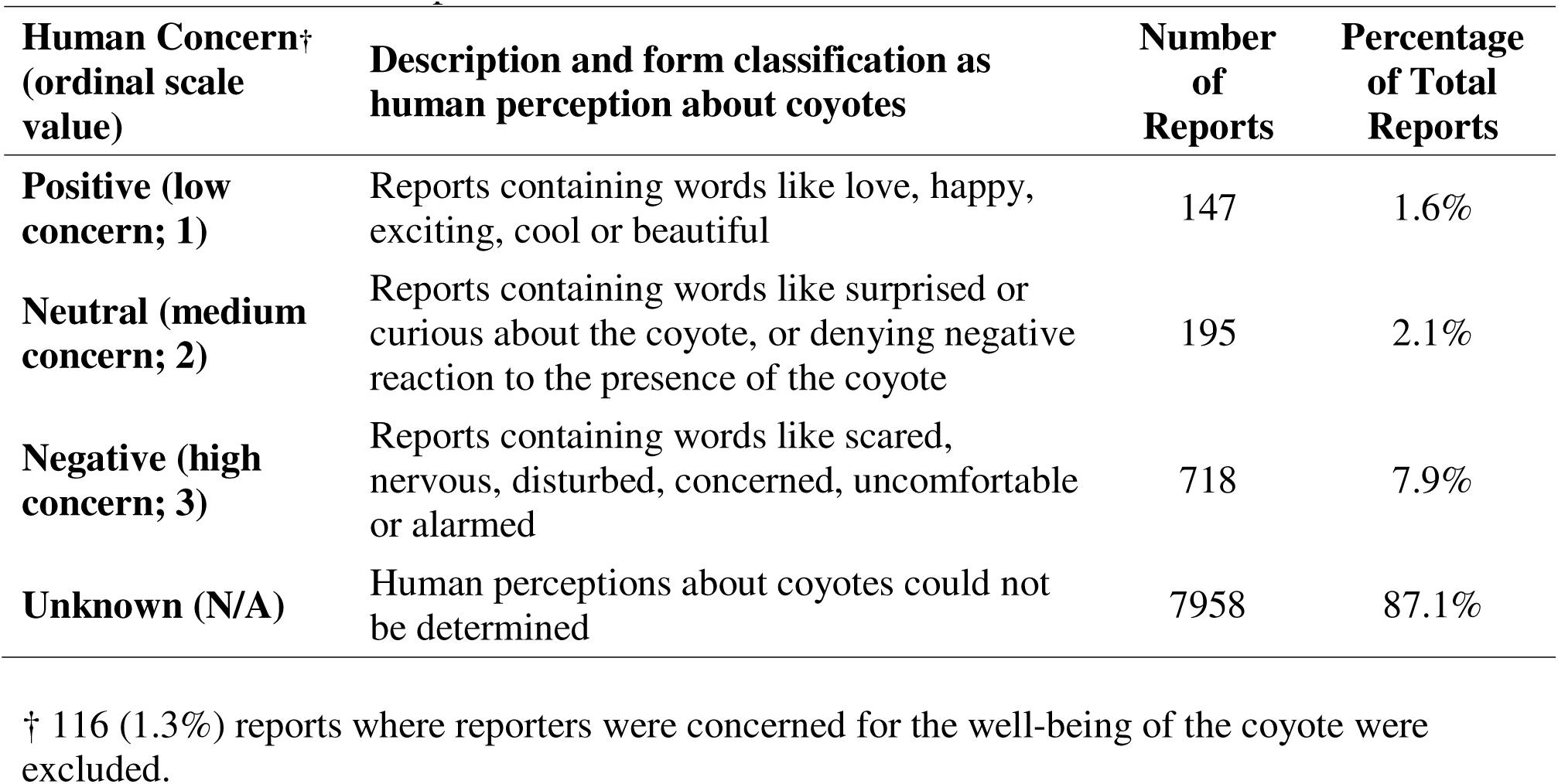
Distribution of reports across human concern ordinal scale values.

| **Human Concern<sup>†</sup>**<br>**(ordinal scale**<br>**value)** | **Description and form classification as**<br>**human perception about coyotes** | **Number**<br>**of**<br>**Reports** | **Percentage**<br>**of Total**<br>**Reports** |
| --- | --- | --- | --- |
| **Positive (low**<br>**concern; 1)** | Reports containing words like love, happy,<br>exciting, cool or beautiful | 147 | 1.6% |
| **Neutral (medium**<br>**concern; 2)** | Reports containing words like surprised or<br>curious about the coyote, or denying negative<br>reaction to the presence of the coyote | 195 | 2.1% |
| **Negative (high**<br>**concern; 3)** | Reports containing words like scared,<br>nervous, disturbed, concerned, uncomfortable<br>or alarmed | 718 | 7.9% |
| **Unknown (N/A)** | Human perceptions about coyotes could not<br>be determined | 7958 | 87.1% |

<sup>†</sup> 116 (1.3%) reports where reporters were concerned for the well-being of the coyote were excluded.

### Spatial and temporal variable collection

To quantify the geospatial setting of each report, we imported report locations into ArcGIS Pro v2.7 (Figure 1). We excluded reports that were located outside of Edmonton city limits or in recently annexed but undeveloped rural land, and we identified our study area by generating a minimum convex polygon around the remaining report locations. Land cover types within our study area were classified using geospatial data from the City of Edmonton Urban Planning Land and Vegetation Inventory (uPLVI) database, a high-resolution database that uses remotely sensed imagery and Softcopy photogrammetry to identify land cover types for urban land use decisions (City of Edmonton 2018). For our study, we binned uPLVI land cover classifications into six land cover types representing various degrees of human development and coyote habitat quality (Table 4) that were comparable to those used in similar studies in the American cities of New York (Weckel et al. 2010), Denver (Poessel et al. 2013), and Atlanta (Mowry et al. 2020).

**Table 4.** Land cover classes representing different degrees of human development and coyote habitat suitability in Edmonton, Alberta as determined from the City of Edmonton Urban Planning Land and Vegetation Inventory (uPLVI) site types.

| Land Cover Class (uPLVI site types) | % Study Area |
| --- | --- |
| 1. Natural<br>(forested, native grass, closed shrub, medial shrub, exposed mineral soil, marsh, treed fen, shrubby fen) | 8.2% |
| 2. Modified Open<br>(annual crops, tame pasture, rough pasture, agriculture hygric tillage site, non-maintained grass/shrub, recent clearing, farmyard/acreage, treed shelterbelt, transplant treed site, nursery/tree farm) | 16.0% |
| 3. Mowed<br>(maintained grass site) | 7.7% |
| 4. Residential<br>(established residential community, residential development site, acreage subdivision) | 42.0% |
| 5. Commercial<br>(established commercial/industrial, commercial/industrial development, aggregates or fill sites, building and/or parking complex, oil and gas field site, transportation surface) | 23.4% |
| 6. Water<br>(anthropogenic water, natural water) | 2.7% |

Because coyote boldness and human concern may be affected by a combination of site-specific conditions (van Bommel et al. 2020) and broader landscape characteristics (Murray et al. 2015b, Wine et al. 2015), we measured land cover at five different scales, within 100, 200, 400, 800 and 1600 m radii of each report. Land cover was calculated as the proportional area of each land cover type within the circular area defined by each scale. Proportional land cover measurements were then transformed using a centered log-ratio transformation to minimize autocorrelation (Quinn et al. 2019). To compare the distribution of reports across different land cover types, we also assigned a single land cover category to each report based on the category with the greatest proportional area within a 100-meter radius of the report.

Building density and road distance have previously been associated with human-coyote encounters (Wine et al. 2015, Drake et al. 2021); therefore, we determined building density based on the proportional area of building footprints within each of the five scales around each report (Statistics Canada 2019). We also measured the distance from each report to the nearest road from the single line street network geospatial database from the City of Edmonton. For road distance, we applied an exponential distance decay function (*e*^−*0.002d*^, where d = meters to the nearest road) to confine values between zero (far from road) and one (on road; Nielsen et al. 2009). All spatial variables were measured in raster format with a 10 × 10-meter cell size.

To support temporal analyses, we measured changes in reporting, coyote boldness, and human concern across years, months, and time of day, as well as across the coyote seasons of breeding (January 1 – April 30), pup rearing (May 1 – August 31), or dispersal (September 1 – December 31). We manually categorized time of day into either day (after sunrise and before sunset) or night (before sunrise or after sunset). Sunrise and sunset times were specific to Edmonton and were adjusted for seasonal variation.

### Statistical methods

We began by categorizing spatial, temporal and contextual patterns in report submissions. For land cover types, we estimated the expected number of reports based on the total proportion of that land cover type within the study area. We then applied Pearson’s chi square test to determine whether reports occurred more or less frequently than expected in each land cover type. To assess how reporting varied over time, we summarized the number of reports in each of the biological coyote seasons for each year from 2012 to 2021, the percentage of reports during each month, and the number of reports from day and nighttime. For each contextual variable we determined the number of reports assigned to categories within each variable.

To identify the best spatial and temporal predictors of coyote boldness and human concern, we used an exploratory modelling framework (Tredennick et al. 2021) based on ordered logistic regression with the *clm* function in the R package ordinal (Christensen 2019). Time of day and contextual variables were strongly correlated with each other (Table A1.1, Appendix 3), so we excluded these variables from our models and examined them separately (see below). We used a pseudo-optimized multiple scale approach (Mcgarigal et al. 2016) to select which of the five measurement scales for each land cover variable and building density fit best for a multivariate model. In brief, this approach involved conducting univariate models for each variable and then retaining the scale with the lowest Akaike’s information criterion value (AIC; Burnham and Anderson 2004; Table A3.2 in Appendix 3), which we did separately for coyote boldness and human concern. If a variable’s best-fit scale did not improve on the AIC of the null model, we excluded that variable from further analyses. We then assessed correlations between the remaining variables using Spearman’s rank correlation coefficient (Tables A3.3 and A3.4 in Appendix 3), and for any pairs of variables where r > 0.6, we removed the variable that produced a higher AIC value in univariate models. All numerical variables were mean centered and scaled with a standard deviation of 1.

For each of coyote boldness and human concern, we constructed global models (Table A3.5 in Appendix 3) that included each of the non-correlated spatial variables, year, and coyote biological season (using breeding season as the reference) as additive effects. We included interaction terms between year and each of the spatial variables to test if temporal changes in the response variables were associated with specific spatial factors in the urban environment. In models of coyote boldness, we also included interaction terms between biological season and each of natural and modified open areas to test for seasonal changes in coyote behaviour that might be associated with denning in these less-developed areas (Dodge and Kashian 2013). Because we were primarily interested in maximizing the explanatory power of our models to explain each of boldness and human concern (Tredennick et al. 2021), we used AIC-based model selection with the *dredge* function from the package MuMIn (Barton 2022) to identify the variables and interactions that were retained in the top models (ΔAIC < 2). To determine whether reports of coyote boldness or human concern had changed over the course of our 10-year dataset, we explored these relationships in more detail using linear regressions predicting the percentage of reports within each of the ordinal scores as a function of year.

We explored the effects of time of day and contextual factors on each of coyote boldness and human concern using Pearson’s chi square tests of independence (Weckel et al. 2010), followed by *post hoc* tests (chisq.posthoc.test.package; Ebbert 2019) to determine which levels of each factor were most strongly associated with boldness or concern. For each of these analyses, we used as a reference category the reports for which the contextual variable could not be determined from reporter comments. We also used chi square tests to determine whether reports that identified bolder coyote behaviour also expressed more human concern, and adjusted alpha values for each residual test with Holm’s correction for multiple comparisons (Macdonald and Gardner 2000). We conducted all statistical analyses in R version 4.1.3 (R Core Team 2022) and considered effects to be significant if 95% confidence intervals did not overlap zero or if p values < 0.05.

## RESULTS

### Reporting patterns

From September 2, 2010, to December 31, 2021, 11,239 reports were submitted on the Edmonton Urban Coyote project website. Of these, we removed 1,722 spam or duplicate reports, 256 reports that were outside of Edmonton city limits, and 127 reports from 2010 and 2011 because of limited reporting in these years. The resulting dataset included 9,134 unique and spatially explicit coyote reports between January 1, 2012, and December 31, 2021. Of the 100 reports that were re-classified to assess classification repeatability, inter-rater agreement for each variable ranged from 85-96% (Table A1.1 in Appendix 1).

Reports were widely distributed across the city and unevenly spread across land cover types (χ^2^_4_= 1,564, p < 0.001; Figure 1). Based on the proportion of each land cover type within our study area, we received more reports than expected in residential (59.1%), mowed grass (12.2%), and natural land cover (10.9 %) areas and fewer than expected in commercial (11.1%) and modified open areas (6.7%). Reporting increased over years, and, within years, was consistently higher in the breeding and dispersal seasons (Figures 2A and 2B). Reports were also more common during the day than at night (Figure 2C). Human activity was discernable in 48.1% of reports and mostly involved walking (19.1%), being in a home or yard (18.4%) or driving (9.1%; Figure 3). Vulnerable individuals (mostly dogs) were present or mentioned in 30.8% of reports, and a subset of those reports identified dogs as leashed (11.7%) or off-leash (9.4%). Most reports involved a single coyote (59.4%), and a subset of reports identified coyote health as healthy (13.9%) or unhealthy (6.0%).

**Figure 2.**
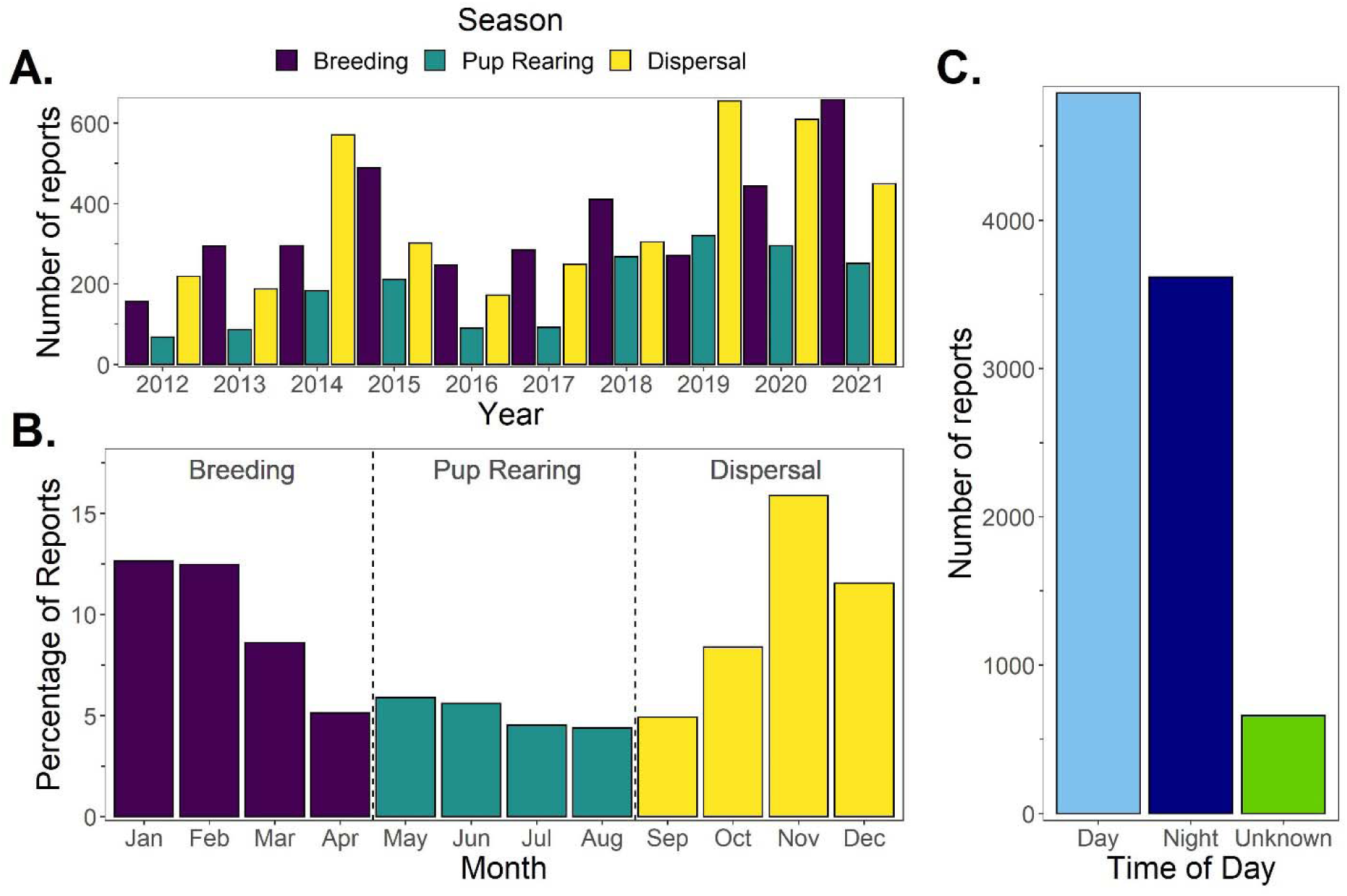
Temporal patterns in coyote reporting across years (A), months (B), coyote seasons (A & B) and time of day (C). Reports were collected through the Edmonton Urban Coyote project website and the date and time of report were submitted by the reporter.

**Figure 3.**
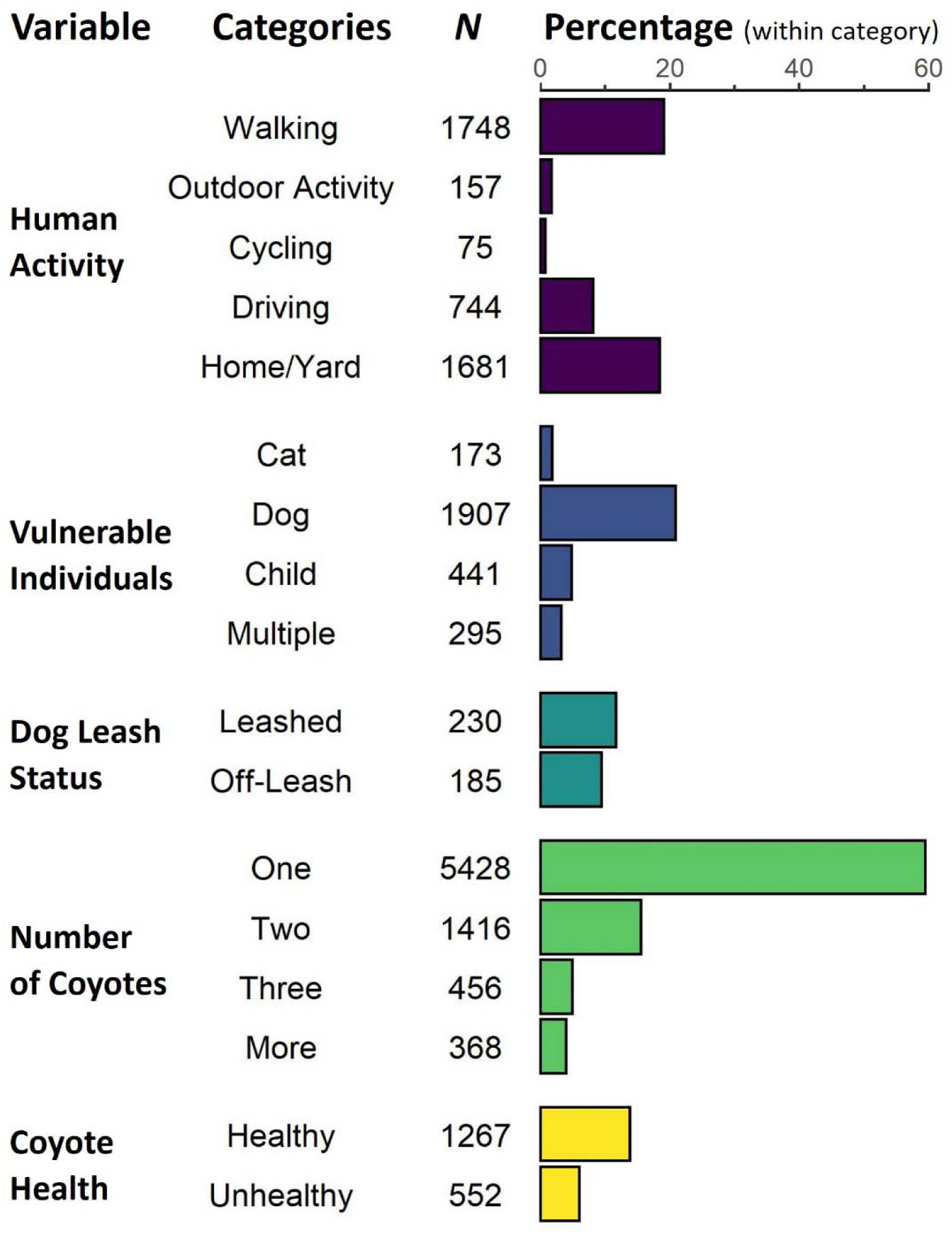
Coyote report distribution across contextual variables. Percentages were calculated based on the total number of reports (*N* = 9,134) with the exception of dog leash status, which is based on the number of reports that mentioned dogs (*N* = 1,958). The remaining (unplotted) reports for each category did not provide relevant information about the contextual variable.

Coyote boldness, as determined from the reported coyote behaviours, was most commonly reported as avoidant or indifferent, followed by bold and aggressive (Table 2). In contrast, human concern, based on the perceptions that people expressed about coyotes, indicated that negativity towards coyotes was much more common than neutral (3.6 times) or positive responses (4.9 times; Table 3). Reports that mentioned physical contact between people or pets and coyotes consisted mostly of dog attacks (*N* = 85), followed by cat depredations (*N* = 50); in only one report did a coyote contact a human while the coyote attempted to take a sled from a child. Among the reports for which both boldness and human concern could be classified, the two variables were significantly related (χ^2^_6_ = 56.3, p < 0.001), with reports of bold or aggressive behaviour being more likely to express negative perceptions of coyotes (Figure 4).

**Figure 4.**
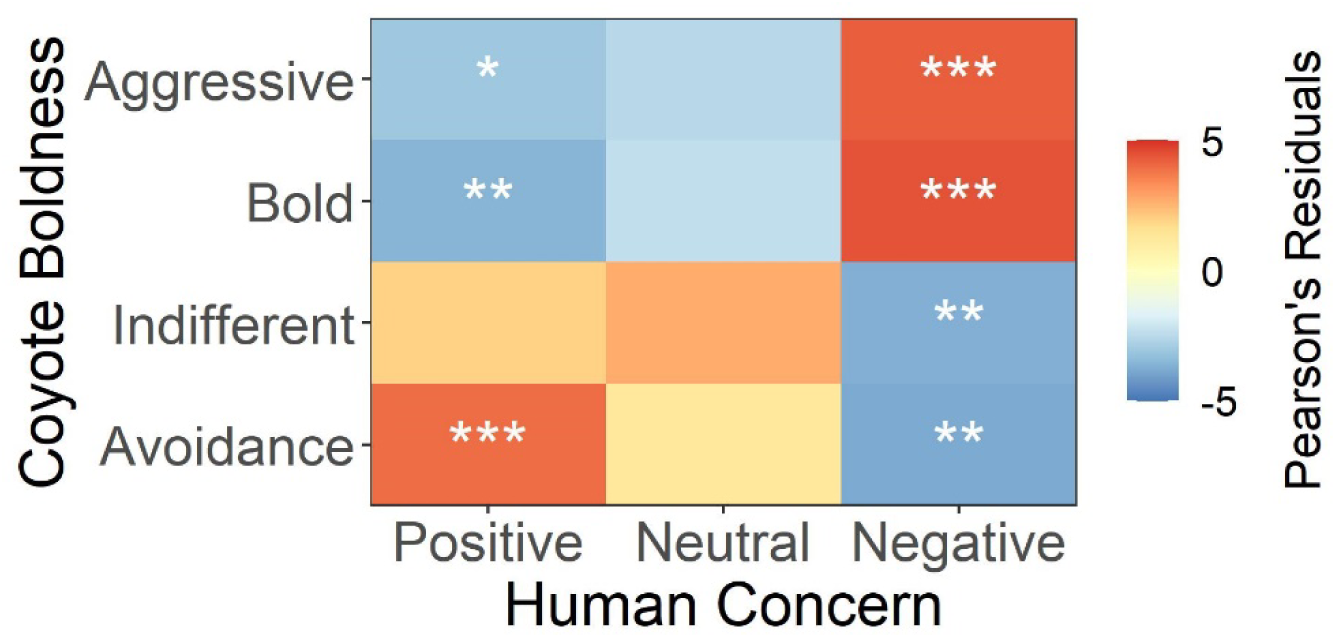
Relationship between coyote boldness, determined from reported coyote behaviour, and human concern about coyotes, determined from participant’s perceptions of coyotes. Colors represent Pearson’s residual values calculated post-hoc from a chi square test, with positive values (red) indicating positive relationships and negative values (blue) indicating negative relationships. Significance is indicated by asterisks (* p < 0.05, ** p <0.01, *** p < 0.001).

### Spatiotemporal predictors of coyote boldness and human concern

Ordered logistic regression analysis revealed a suite of spatial and temporal variables that explained each of coyote boldness and human concern (Figure 5). We present results from only the top models for these response variables (Figure 5) because there was little variation among coefficient and confidence estimates within the full set of top-ranked models (ΔAIC_c_ < 2; Tables A3.6 and A3.7 in Appendix 3). The top model predicting coyote boldness indicated that the log odds likelihood of bolder behaviour was higher during the pup rearing season and in areas with higher proportions of mowed land cover (within a 100 m radius), but lower closer to roads and in areas with greater building density (within 200 m). The significant interaction term in this model indicated that boldness was higher during the pup rearing season especially in areas with more modified open land cover (400 m buffer size, Figure A3.1 in Appendix 3). The top model for human concern indicated that a higher likelihood of human concern was associated with increases in the proportion of residential area (within 800 m) and modified open land cover (within 1600 m), as well as during the dispersal season (Figure 5)

**Figure 5.**
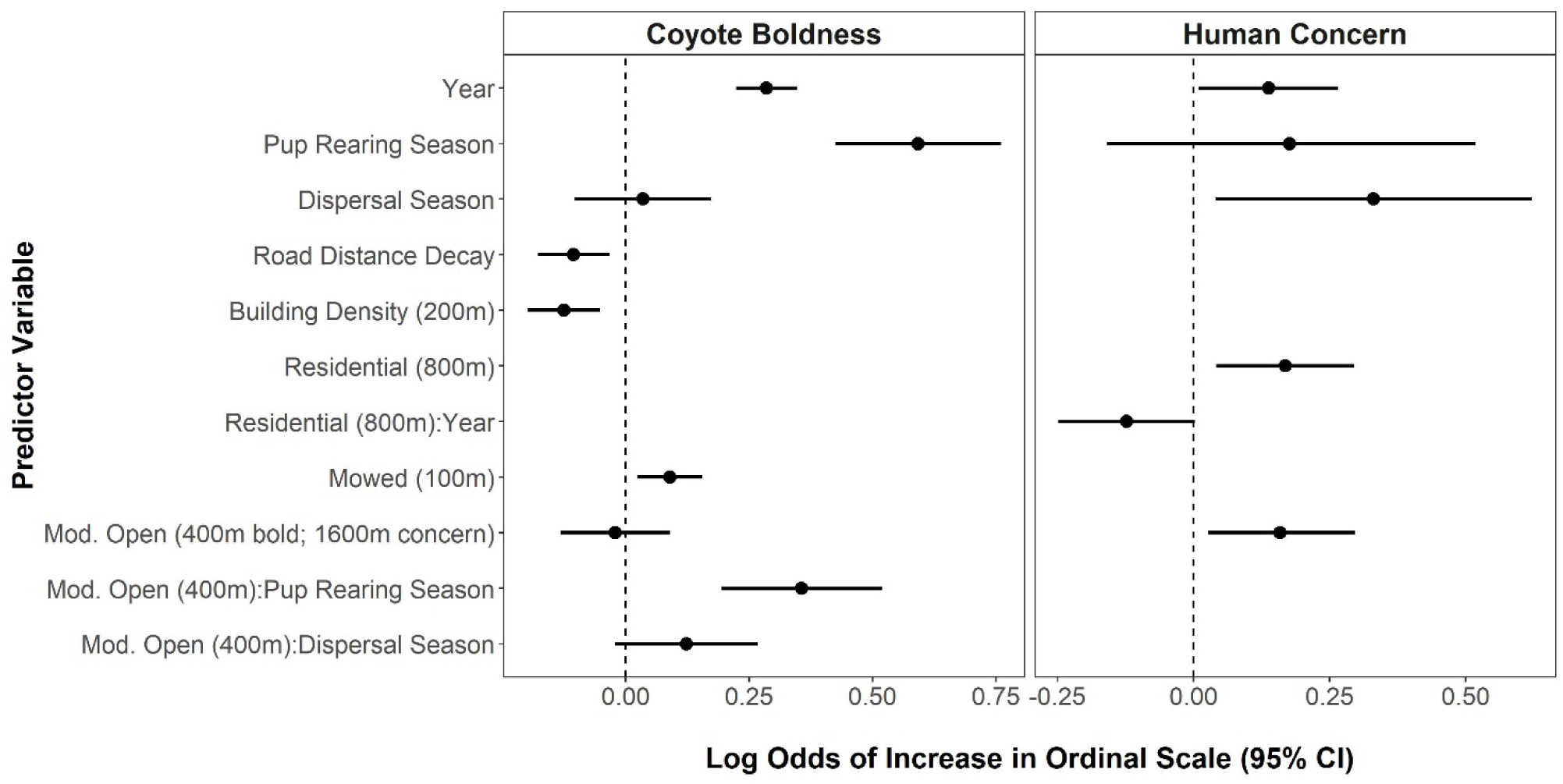
Log odds coefficient values and 95% confidence intervals for the explanatory variables retained in the top ordered logistic regression models (lowest AIC_c_) for coyote boldness and human concern of coyotes. The full set of variables included in each global model is available in Table 5 of Appendix 3. Positive values indicate that the variable causes a higher likelihood of conflict-associated coyote behaviour or human concern, while negative values suggest reduced likelihoods.

Models for both coyote boldness and human concern indicated a significant increase in the likelihood of human-coyote conflict over years (Figure 5). None of the interaction terms with year were significant when we considered only the single best model for each response variable (Table A3.6, Table A3.7 in Appendix 3); however, for models predicting human concern, the negative interaction term between residential area and year was retained in 19 of 20 top models and was significant in 13 of these. This interaction indicated that concern was generally higher in residential areas in early years, but increasing levels of concern in non-residential areas reduced the magnitude of the effect of residential area over time (Figure A3.2 in Appendix 3). While several other variables and interaction terms appeared in some of the top models (ΔAIC_c_ < 2) for boldness and concern, their effects were not significant in any of these (Tables A3.6 and A3.7 in Appendix 3).

We examined temporal changes in greater detail by evaluating the percentage of reports within each of the ordinal scores for each year (Figure 6). Specifically, the percentage of reports describing bold behaviour increased significantly (β = 2.19, p < 0.001) while avoidance behaviour decreased (β = −1.82, p < 0.001), though there were no differences in the percentage of reports describing indifferent (β = −0.71, p = 0.21) and aggressive behaviour (β = 0.24, p = 0.16). Similarly, negative perceptions about coyotes became more common over years (β = 1.07, p = 0.072) and positive perceptions became less common (β = −1.07, p = 0.005), with no change in the proportion of neutral perceptions (β = 0.002, p = 0.997).

**Figure 6.**
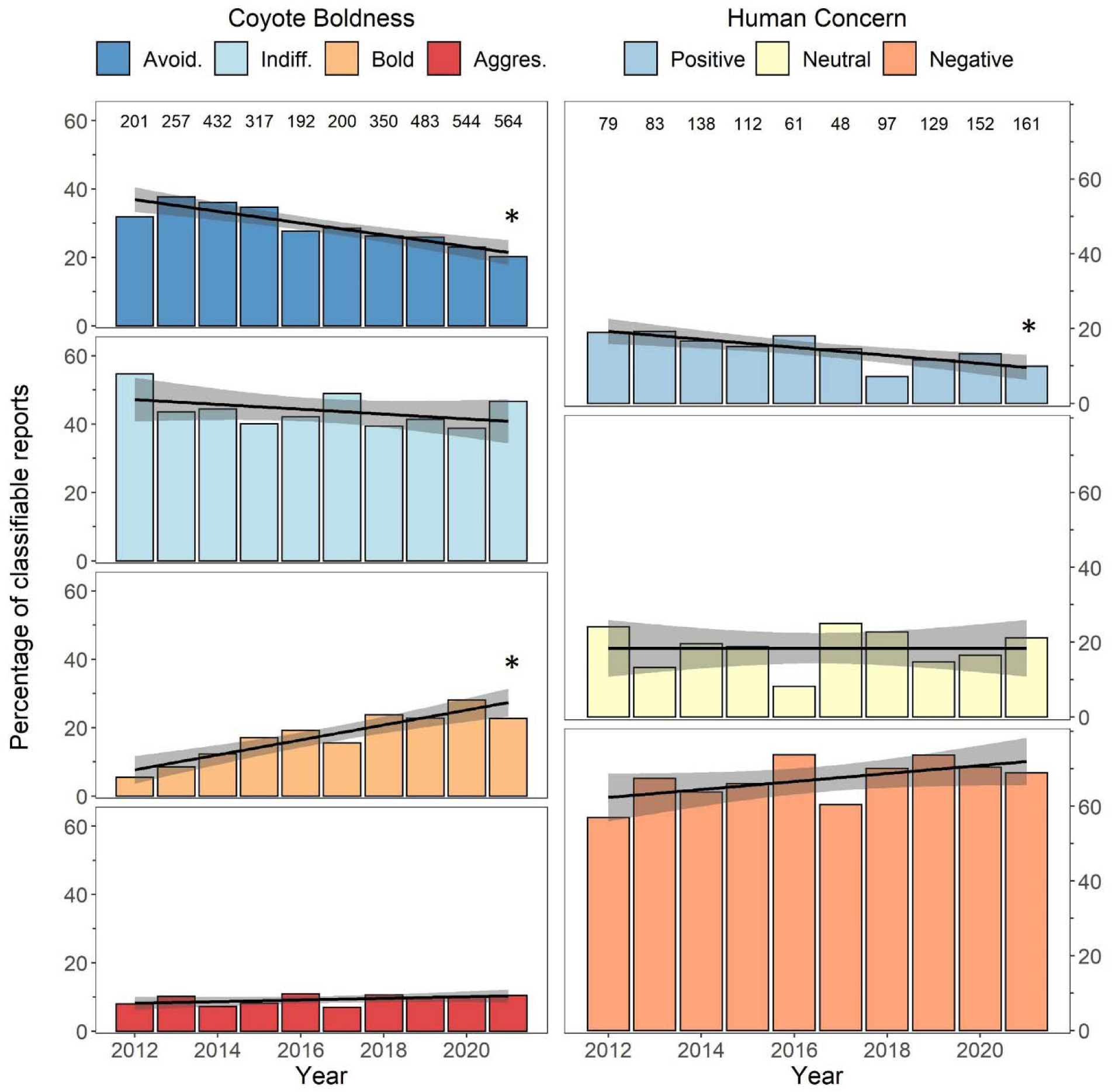
Long-term (10 year) trends in coyote boldness and human concern of coyotes as indicated by the percentage of reports in each of the boldness or concern categories. Reports were collected through the Edmonton Urban Coyote project website, and boldness and human concern were scored on ordinal scales using predetermined criteria. Numbers at the top of each chart denote the total number of reports for each year for which an ordinal score could be assigned. Linear trends are shown with 95% confidence intervals shaded in grey and significant trends are indicated by asterisks (p < 0.05).

Analysis of diel patterns in coyote boldness showed that indifferent behaviour was significantly more common during the day and avoidance behaviour was significantly more common at night (χ^2^_2_ = 30.1, P < 0.001; Figure A2.1 in Appendix 2). However, human concern did not differ between day and night (χ^2^_2_ = 1.09, P = 0.58).

### Contextual influences on boldness and concern

All five contextual variables were significantly related to coyote boldness (Figure 7, Table A2.3 in Appendix 2). Bold behaviour was described more frequently when reporters were walking, when dogs were mentioned, and when two or three coyotes were present. Aggressive coyote behaviour was reported more frequently than expected when cats or dogs were mentioned, when dogs were off-leash, and when two or more than three coyotes were observed. The least threatening coyote behaviours, avoidance and indifference, occurred mostly in reports when people were driving, cycling, or in their home or yard, when only one coyote was observed, and when coyotes were perceived as healthy.

**Figure 7:**
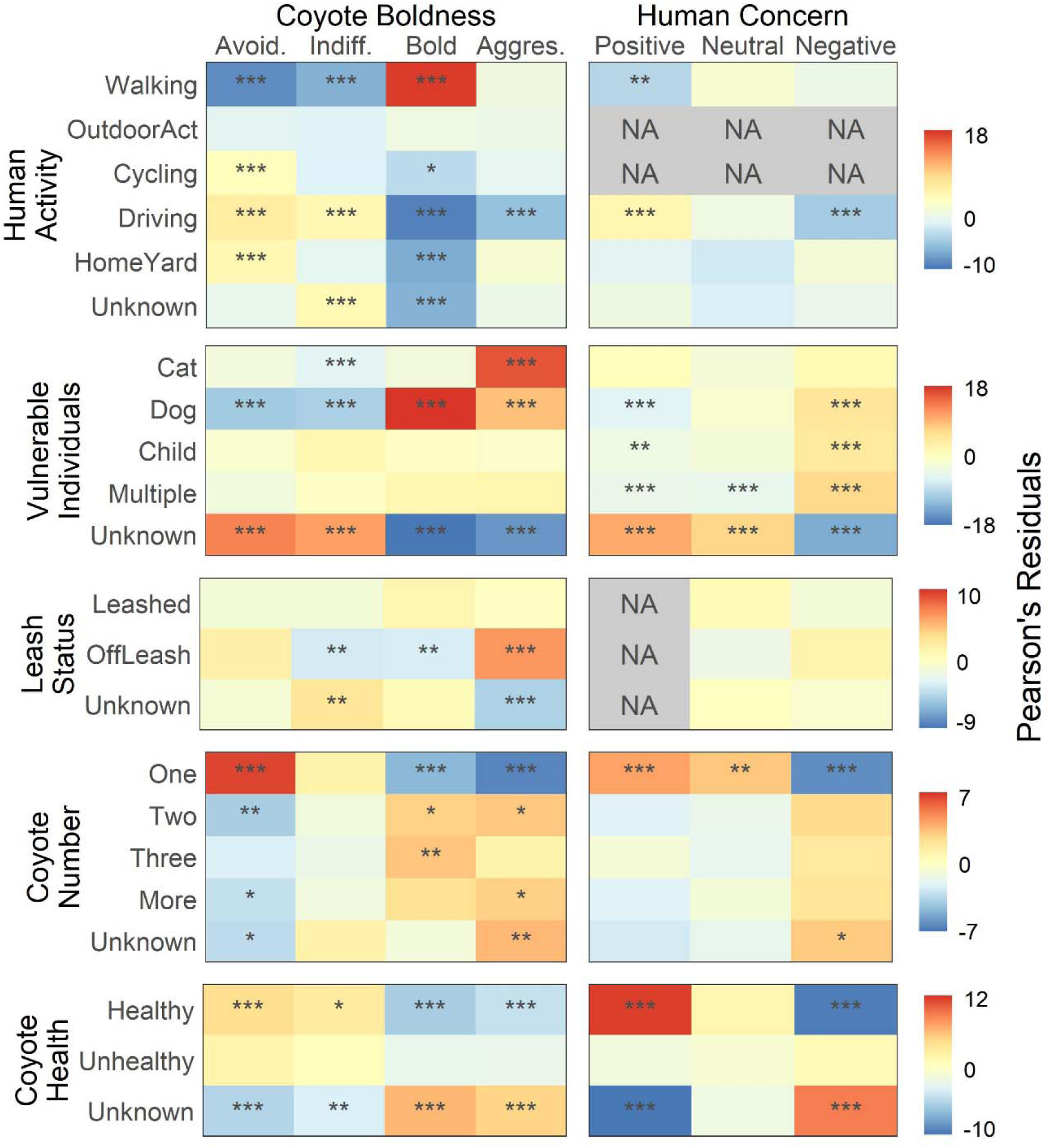
Relationship between each of coyote boldness and human concern of coyotes with contextual independent variables (human activity, the presence or mention of vulnerable individuals, dog leash status, coyote number and coyote health). Colors represent Pearson’s residual values calculated post-hoc from statistically significant chi-square tests of independence, with positive values (red) indicating positive relationships and negative values (blue) indicating negative relationships. P values were adjusted for multiple comparisons with Holm’s correction and significance is indicated by asterisks (* p < 0.05, ** p <0.01, *** p < 0.001). Grey boxes (NA) indicate comparisons for which insufficient reports were available to allow for robust chi square tests (< 5 expected reports).

Most contextual variables were also related to human concern, demonstrating that they were important factors affecting the perceived risk presented by coyotes (Figure 7, Table A2.3 in Appendix 2). Concern was more frequently reported when dogs, children, or multiple vulnerable individuals were mentioned; conversely, reports that didn’t mention any vulnerable individuals expressed less concern. Perceptions were more likely to be positive when only one coyote was observed and when the coyote(s) were described as healthy.

## DISCUSSION

Human-coyote conflict is increasing in urban areas throughout North America (White and Gehrt 2009, Baker and Timm 2017), creating a need to better understand the predictors of conflict so that they can inform proactive management strategies. We used data from a 10-year database of citizen reports to explore which spatial, temporal and contextual predictors best explained the degree of boldness in descriptions of coyote behaviours and concern in perceptions of coyotes expressed by reporters. We found that descriptions of coyote boldness increased: in areas with more mowed grass, with lower building density, and as distance to roads increased; over time and in the pup rearing season; and when reporters were walking, mentioned a cat or dog, and when more coyotes were present. Human concern was greater: in areas with higher proportions of residential and modified open land cover; over time and during the dispersal season; and when reporters mentioned vulnerable individuals. Both boldness and human concern increased significantly from 2012 to 2021.

### Spatiotemporal patterns in coyote boldness

We found that coyote boldness was higher in less-developed areas that were mowed or otherwise not naturally vegetated, similar to what was described for open relative to natural areas in Denver, Colorado (Poessel et al. 2013) and in managed clearings in North Carolina (Wine et al. 2015). Collectively, these results suggest that conflict-indicative coyote behaviour is most prevalent in spaces that are at the interface of natural and developed urban areas. Similar interfaces between peri-urban and rural or rural and wildland areas frequently concentrate human-wildlife conflicts in many other species (König et al. 2020). The pattern we observed may arise because coyotes in open areas are visible at greater distances, and may thus appear to be bolder; alternatively, bolder animals may be more likely to occupy areas with less vegetation cover, as has been reported for brown bears (*Ursus arctos*; Bombieri et al. 2021). This behaviour might be expected of coyotes owing to their evolution in the arid southwest of the North American continent (Hody and Kays 2018). Our spatial variables were most explanatory of boldness when measured at smaller spatial scales (≤ 400 m radii from reports; Table A3.2 in Appendix 3), suggesting that boldness is driven by site-specific factors, like proximity to vegetation cover, territorial boundaries, or den sites.

Seasonally, boldness was significantly more likely during the summer pup rearing season, which is also consistent with other studies (White and Gehrt 2009, Lukasik and Alexander 2011). The fact that fewer reports were submitted during this period, despite a time of greater outdoor activity by people in our study area, suggests that coyotes avoid humans and pets during pup rearing, but behave more aggressively when interactions occur. Aggression by coyotes during the pup rearing season presumably reflects defence of pups from perceived threats posed by humans or dogs (Bombieri et al. 2018). Many reports described coyotes rushing out of cover to bite large dogs on their hamstrings, suggestive of defensive behaviour. There were also more reports of cat depredations during the pup rearing season (29 of 50 total), probably caused by a combination of coyotes seeking food for their pups and generally greater numbers of free-roaming cats in the summer that might, in turn, be hunting naïve, young birds and rodents (Nation and St. Clair 2019).

Boldness during the pup rearing season was particularly associated with modified open areas, which may have been associated with denning by coyotes in these areas. Dens in modified open areas like utility corridors, meadows within the city’s ravine system, or agricultural land near and within the city have less vegetative cover than dens in natural areas, which might reduce opportunities for avoidant behaviour and increase defensive aggression. Indeed, reports from natural areas were more likely to describe avoidance (28.9%) than aggression (12.9 %) relative to reports from modified open areas (20.3% avoidance and 20.7% aggression; Table A2.1 in Appendix 2). An alternative hypothesis is that coyotes denning in more disturbed modified open areas may be more prone to boldness because of repeated exposure and thus habituation to humans and their pets in these areas (Young et al. 2019). In either case, our study shows that coyote behaviour in modified open areas during the pup rearing season may often present a risk to the safety of humans and their pets.

We additionally found that coyote boldness increased over the 10-year reporting period, potentially explaining the mechanism for rising coyote attacks on people in North America (White and Gehrt 2009, Baker and Timm 2017). Both patterns may reflect the greater boldness of urban, relative to rural coyotes that others have reported and attributed to reduced persecution by people, repeated benign interactions with humans, and access to anthropogenic food (Breck et al. 2019, Young et al. 2019, Brooks et al. 2020). There is evidence that coyote boldness towards humans is passed from parents to offspring (Schell et al. 2018) and increases with greater exposure to people (Young et al. 2019), both of which could increase coyote boldness over time to accelerate boldness-driven conflict. Separate from these interactions with people, higher coyote population density within cities may lead to intraspecific competition that favours bolder individuals (Bateman and Fleming 2012). Our top ordinal regression models did not include any interactions between spatial variables and year, suggesting that these changes in boldness were relatively consistent across the urban environment. Despite increases in boldness over time, we did not find a similar increase in aggressive behaviour, possibly because the most aggressive individuals were targeted for removal by city managers, which is an effective means of reducing conflict in the short term (Breck et al. 2017).

### Spatiotemporal patterns in human concern

We classified human perceptions of coyotes described in reports as positive, neutral, and negative to create a metric of human concern that increased with the amount of residential area within 800 m of the report. This observation is similar to previous findings that people are less tolerant of coyotes near their homes despite being generally tolerant of coyotes in cities (Bonnell and Breck 2017, Drake et al. 2020). A similar pattern occurred for cougars (*Puma concolor*) at a rural-wildland interface in Alberta (Knopff et al. 2016). Such effects show how human concern may not align with the actual risk of a coyote behaving boldly or aggressively: concern was higher in residential areas, but boldness was negatively associated with building density and road proximity, which were correlated with residential area in our study (Table A3.3 in Appendix 3).

Human concern about coyotes was also higher in areas with more modified open land cover, where bold interactions are more likely during the pup rearing season. This land cover may also be disproportionally responsible for the positive correlation we found between reports that described bolder coyotes and greater human concern. Such a correlation might be amplified if people gain awareness over time of the bold or aggressive coyote interactions that are more common in those areas, which has previously been shown to increase the risk people perceive from coyotes (Sponarski et al. 2018, Draheim et al. 2019). People may respond most to the visibility of coyotes, which is likely highest during the fall dispersal season when our database received more reports and higher levels of concern. Interestingly, human concern correlated with land cover predictors measured at larger scales (≥ 800 m radii; Table A3.2 in Appendix 3) than did the measures of boldness derived from descriptions of coyote behaviour. This difference may reveal that coyote behaviour and human perceptions are affected by spatial variables at different scales, or, simply, that coyote behaviour is more readily described for animals that are nearby.

Like boldness, human concern about coyotes increased over the 10-year reporting period (Figure 5, 6), and our study demonstrates that these perceptual changes may be quite nuanced across the urban landscape. For example, concern was initially higher in areas with more residential land cover, but this effect was reduced over time due to growing human concern across most other land cover types. While some studies have predicted that humans will habituate to the presence of urban coyotes over time (Lawrence and Krausman 2011, Jackman and Rutberg 2015), as has been reported in Alaskan brown bears (Smith et al. 2005), our findings support suggestions by others that this does not occur with carnivores if people fear for their safety (Williams et al. 2002, Kaltenborn et al. 2006). Expecting that human concern will naturally decrease as coyotes become more prevalent in cities may not be an effective strategy to facilitate coexistence, because public concern about any carnivore may typically increase with increasing carnivore prevalence, negative interactions, or if conflict is emphasized by the media (Lute and Carter 2020).

### Contextual factors affect coyote boldness and human concern

Coyote boldness and human concern were each predicted by several contextual variables. Reports describing the presence or mention of vulnerable individuals exhibited higher scores for both coyote boldness and human concern. The mention of dogs or cats were associated with descriptions of bolder and more aggressive coyote behaviour, supporting the findings of others that human-coyote interactions involving pets are more likely to cause conflict (Poessel et al. 2013, Baker and Timm 2017). Coyotes were described as “aggressive” less often when dogs were leashed (14.6%) compared to when they were described as off leash (32.5%; Table A2.1 in Appendix 2), but bold behaviour was more common when dogs were leashed (39.2%) than off leash (22.3%). This result suggests that leashed dogs may still engender coyote behaviour that is associated with conflict even if they are less likely to be attacked than when they are off leash. Many of these interactions during the pup-rearing season may have involved escorting behaviour, wherein coyotes do not attack, but follow people with leashed dogs out of areas near their den sites. We did not find a significant relationship between the presence or mention of children and coyote boldness or aggression. Nonetheless, human concern was significantly higher when reports mentioned dogs, children, or multiple vulnerable individuals, perhaps because occasional coyote attacks on children are well-publicized and can lead to serious injuries (Carbyn 1989, White and Gehrt 2009, Alexander and Quinn 2011; Baker and Timm 2017). A higher perceived risk of coyote attacks on pets and children likely reduces tolerance for coyotes in cities (Draheim et al. 2019).

With respect to other contextual variables, coyotes were described as being bolder when they were observed with other coyotes and when people were walking, whereas expressions of human concern were lower when people were driving or when only a single coyote was mentioned. Although coyotes in poor health may be more conflict-prone (Murray et al. 2015b), we found no evidence that people found unhealthy animals to be bolder or feared them more. However, coyotes that were perceived to be healthy were more likely to be described as avoidant or indifferent. It could be that reporters were less likely to notice and characterise a coyote’s health in encounters where coyotes were behaving boldly or aggressively.

### Limitations

Our study had several limitations. First, reports were collected non-randomly and non-independently, which introduces several potential biases inherent to community reporting databases (Poessel et al. 2013, Sullivan et al. 2014). These biases include greater tendencies for repeat reporting by some residents with particularly strong views about coyotes, uneven advertising of the reporting database across neighborhoods or over years, potentially higher reporting from affluent neighborhoods with higher education levels (Wine et al. 2015, Mowry et al. 2020), and varying visibility of coyotes across seasons, time of day or land cover types due to differences in vegetative cover, human activity and daylight (Quinn 1995, Poessel et al. 2013). We attempted to mitigate these effects by focusing on measures of coyote boldness and human concern, rather than spatiotemporal influences on the number or distribution of reports. Additionally, we attempted to overcome spatial and temporal autocorrelation in the reports that could contribute to Type 1 statistical errors by restricting analyses to those with large sample sizes and verifying modelling results with chi-square tests (Table A2.3 in Appendix 2). Despite these precautions, our post-hoc method of quantifying coyote boldness and human concern from a community reporting database cannot be compared to empirical behavioural observations of animals (e.g., Breck et al. 2019) or randomized public surveys (e.g., Drake et al. 2020).

### Management implications

Despite some limitations, our findings support the increased implementation of several management actions in our study area and elsewhere by identifying areas, seasons and contexts that are associated with higher rates of human-conflict. Spatially, coyote boldness was generally higher in less developed and open areas (Poessel et al. 2013, Wine et al. 2015) where managers might emphasize public education and protection of pets via leashing and fenced areas (Draheim et al. 2019). Managers could also target these areas for initiatives to train community members to haze coyotes that behave boldly (Bonnell and Breck 2017). Both awareness campaigns and community-based hazing programs might target these conflict-prone areas in and near residential neighborhoods, where human concern of coyotes is highest, especially during the breeding season to prevent denning in these areas and proactively limit negative interactions (Bonnell and Breck 2017). Further management actions to improve coexistence should include attractant management in residential areas to reduce coyote attraction to anthropogenic food sources (Murray and St. Clair 2017), and even the targeted removal of particularly aggressive or food conditioned individuals (Breck et al. 2017). The removal of such individuals, although controversial (Breck et al. 2017. Draheim et al. 2019), may be needed to prevent attacks on pets or people that can generate substantial media attention and erode public tolerance of coyotes (Alexander and Quinn 2011, Draheim et al. 2019). We also suggest that managers should address contextual variables while acknowledging the different scales of our findings to target bold coyote behaviour in localized areas (e.g., areas where people walk dogs off leash or schoolyards), while addressing human concern of coyotes at larger scales (e.g., neighborhoods). Finally, we found a rise in both coyote boldness and human concern from 2012 to 2021, and, as such, expect that conflict may continue to rise without more effective and extensive implementation of the aforementioned strategies to reduce negative human-coyote interactions (Kaltenborn et al. 2006, Lute and Carter 2020), facilitate positive wildlife experiences (Kretser et al. 2009), and increase public knowledge about risk reduction (Riley and Decker 2000). Future studies using community reporting databases like ours will provide even greater insight to the spatiotemporal and contextual predictors of conflict with coyotes, and will further facilitate human-coyote coexistence in cities across North America.

## Acknowledgements

We respectfully acknowledge that this work was conducted on Treaty 6 territory, a traditional gathering place for diverse Indigenous peoples including the Cree, Blackfoot, Métis, Nakota Sioux, Iroquois, Dene, Ojibway/Saulteaux/Anishinaabe, Inuit, and many others. We are grateful to Tobias Tan for building the reporting site, and the thousands of Edmontonians who submitted reports of urban coyotes with detailed descriptions that made our study possible. =We thank Cassondra Stevenson for assistance with geospatial analyses, and Sage Raymond and Arya Horon for providing valuable feedback on this manuscript. We deeply appreciate the volunteers and undergraduate students who donated their time to help classify coyote reports (Abby Keller, Amy Malo, Asma Hamid, Arya Horon, Allison Cain, Caley Campkin, Cleo Randall, Donovan Currie, Danika Wack, Emilie Torwalt, Elizabeth Blanchette, Gabrielle Lajeunesse, Hailey Dunsire, Jessica Butts, Jonathan Wild, Kelsey Fleming, Khoi Nguyen, Maria Diaz, Matthew Elphick, Muskaan Tiwari, Osa Campbell, Rachel Godinho, Sage Raymond, Sydney Enns, Sofia Guest, Stephen Shikaze, Tawnee Dupuis, and Vala Ingolfsson). Funding for this study was provided by a Discovery Grand from the Natural Science and Engineering Research Council of Canada and a research fellowship from the Faculty of Science, University of Alberta to CCSC.

