## Appendix 1 for "A ten-year community reporting database reveals rising coyote boldness and associated human concern in Edmonton, Canada"

### APPENDIX 1. COYOTE REPORT CLASSIFICATION

**Figure A1.1.** The report classification form used to extract information from coyote reports. Volunteers read each report and completed this form.

#### Coyote Report Classification

This is only to be used for classifying the reports that are no longer training reports (real reports only!)

* Required

1. Report ID (NOT the excel row number) *

    ______________________________

2. Number of Coyotes *

    *Mark only one oval.*

    - ◯ One (e.g., name 'a coyote' or single, lone)
    - ◯ Two (e.g., pair, couple)
    - ◯ Three (including a few)
    - ◯ More
    - ◯ Unknown

3. Coyote young are named (including dens)

    *Check all that apply.*

    - ☐ Yes

4. Human activity

*Mark only one oval.*

- ◯ Walking
- ◯ Driving
- ◯ In home or yard
- ◯ Cycling
- ◯ Designated outdoor activity: jogging, golfing, hiking
- ◯ Unknown
- ◯ Other: ____________________

5. Vulnerable individual present or implied *

*Mark only one oval.*

- ◯ Child
- ◯ Dog
- ◯ Cat
- ◯ Multiple (e.g., pets and children)
- ◯ Unknown
- ◯ Other: ____________________

6. If a dog was present, was it: *

*Mark only one oval.*

- ◯ Leashed
- ◯ Off-leash
- ◯ In home / yard
- ◯ Unknown
- ◯ Other: ____________________

7. Did the person try hazing the coyote? *

*Mark only one oval.*

- Yes
- No
- Unknown

8. Coyote response to people *

*Mark only one oval.*

- 0. Did not / could not see the people
- 1. Ran away
- 2. Walked away
- 3. Did not appear to notice or care about people
- 4. Watched the person
- 5. Howled at the person
- 6. Followed or stalked pets or people
- 7. Approached the person
- 8. Chased or charged pets or people
- 9. Made physical contact with pets or people
- Unknown
- Other: ______________________________

9. Human Perception *

*Mark only one oval.*

- Reporter expresses negative perception of coyote or emotion
- Reporter expresses neutral perception or emotion
- Reporter expresses positive perception or emotion
- Reporter expresses concern for coyote
- Unable to determine

10. Coyote health is mentioned *

*Mark only one oval.*

- ◯ Healthy (e.g., Healthy, strong, beautiful, or majestic)
- ◯ Unhealthy (e.g., Injured, limping, hobbling, hurt, unhealthy, mange, mangy, scruffy, or wounded)
- ◯ Unknown
- ◯ Other: ______________________

11. Classifier ID (first initial and last name) *

______________________

12. Additional Comments

______________________
______________________
______________________
______________________
______________________

13. Report Classification should be reviewed

*Check all that apply.*

- ☐ Yes

This content is neither created nor endorsed by Google.

Google Forms

**Text A1.1.** The protocol used to train volunteers and inform their use of the report classification form.

*Volunteer Training Protocol*

**1)** Initial Training: Volunteers will be sent 30 reports to classify, following the classification protocol outlined below. Each of these reports has been previously classified by Author. Once volunteers have classified the 30 reports, they will notify Author and a virtual meeting will be scheduled to go over the reports and examine where answers differ from author's classifications. Author will explain why he classified the report differently to help the volunteer adjust their coding to minimize variability.

**2)** Secondary Training: Volunteers will then classify another 30 practice reports, and compare their answers to the classification done by Author (on their own time). If necessary, they will contact Author with any questions.

**3)** Report Classification: Once volunteers feel confident in their classification ability (steps 1 and 2 can be repeated as needed) they will be sent 100 reports at a time to be classified. All reports shared with volunteers will have all personal identifiers (name, contact, location) removed.

*Google Form Report Classification Protocol*

The descriptions below are stepwise guidelines to help volunteers classify reports. Please direct any questions to Author. Remember, it is important to avoid making inferences when classifying the reports.

**1)** Report ID
- Record the unique ID of the report being classified (not the spreadsheet case/row)

**2)** Number of coyotes
- Select the bullet point that best matches the number of coyotes defined in the report

**3)** Coyote young are named
- Select the box only if coyote young (young/pups/babies) or dens are named explicitly (NOT small/little/tiny)

**4)** Human activity
- Select the box that best fits the activity described in the report. If no activity is explicitly mentioned (for example, if a report says "saw coyote in alley") select *Unknown.*

**5)** Vulnerable individual present or implied
- The vulnerable individual does not need to be explicitly present, if reports say "near a school" or "worried because children play in this area," select *Child.*

**6)** If a dog was present, was it:
- Select *Leashed, Off-leash* or *In home / yard* if explicitly stated or if dog is mentioned, if dog is not mentioned or is not explicitly stated, select unknown.

**7)** Did the person try hazing the coyote?
- Select *Yes* only if the person explicitly mentions trying to haze the coyote by chasing, shouting, kicking, throwing things or honking at the coyote.
- Select *No* if the person explicitly attempts to avoid interaction with the coyote by walking away, running away or standing still. Also select *No* if the person was in a situation where hazing was not possible, for example if the coyote did not notice the person, if they saw the coyote on security footage, or viewed it from a distance.
- Select *Unknown* for all other situations.

**8)** Coyote response to people
- In this section, it is important to read the entire report then select the highest option that the report indicates. For example, if the coyote watched the person then approached them

and bit their dog, select *Made physical contact with pets or people*, NOT *Watched the person* or *Approached the person.*

  - Select *Did not / could not see the people* if the coyote is described as being seen from a distance, or from indoors, or if the coyote is explicitly described as not seeing the person. If the coyote is described as preoccupied (hunting, sniffing around) without giving any further details, select *Did not / could not see the people*
  - Select *Ran away* if the coyote explicitly avoided human interaction by running away. A report saying they saw a coyote running across a street should not be classified as *Ran away*. If the coyote is running away from an off leash dog, make note of this in the comment
  - Select *Walked away* if the coyote explicitly avoided human interaction by walking away. A report saying they saw a coyote walking across a street should not be classified as *Walked away.*
  - Select *Did not appear to notice or care about people* if the coyote is described as being indifferent to people.
  - Select *Watched the person* only if the coyote is explicitly described as watching the person (looking at them) without approaching them
  - Select *Howled at the person* only if the coyote is howling/yipping at the person directly. Reports that are auditory descriptions of coyotes howling outside at night in the ravine or neighbourhood should not be classified as *Howled at the person.*
  - Select *Followed or stalked pets or people* only if the coyote is described as following/escorting/stalking the people without attempting to approach them.
  - Select *Approached the person* only if the coyote is described as approaching/nearing/sneaking up on the person.
  - Select *Chased or charged pets or people* if the coyote lunges/chases/bites at the person, and does not make contact because it is hazed by the person or decides not to make contact for another unknown reason
  - Select *Made physical contact with pets or people* only if the coyote bites or is kicked/punched/bitten by a person or pet.
  - Select *Unknown* if the report is too vague to infer any of the other responses. For example "Saw coyote in field" or "Coyote ran/walked across street" would be classified as *Unknown*

**9)** Human Perception

- *Reporter expressed negative perception of coyote or emotion* should be selected if the report uses words such as terrified, scared, uncomfortable, nervous, worried, frightened, disturbing, fear, concern, or alarm.
- *Reporter expresses neutral perception or emotion* should be selected if the report uses words such as surprised, curious or denied a negative reaction, such as "wasn't scared."
- *Reporter expresses positive perception or emotion* should be selected if the report uses words such as love, happy, exciting, cool, and beautiful.
- *Reporter expresses concern for coyote* should be selected only if the person explicitly expresses worry or concern for the coyote's well-being.
- *Unable to determine* should be selected for all reports where the reporter does not describe the coyote using one of the words above, or does not address their emotional response to the coyote sighting/encounter using one of the words above.

**10)** Coyote health is mentioned

- Select the box that best describes the health of the coyote if described in the report.

**11)** Additional comments

- Please record any additional comments you may have

**12)** Classifier ID (first initial and last name)

- The first initial and last name of the person who classified the report (ex: A. Author)

**13)** Reporter comments should be reviewed

- Select only if you desire for your comments to be reviewed by Author. You may also or text XXX-XXX-XXXX with any questions that may come up.

**Table A1.1.** Inter-observer agreement between report classifiers.

| **Variable** | **Inter-rater agreement†** |
|---|---|
| Coyote boldness | 85 % |
| Human concern | 96 % |
| Human activity | 87 % |
| Vulnerable individual | 97 % |
| Dog leash status | 85 % |
| Number of coyotes observed | 92 % |
| Coyote health | 89 % |

† Percentage of reports out of 100 classified the same between two classifiers
