## Appendix 2 for "A ten-year community reporting database reveals rising coyote boldness and associated human concern in Edmonton, Canada"

### APPENDIX 2. COYOTE BOLDNESS AND HUMAN CONCERN ACROSS VARIABLES

**Table A2.1.** Distribution of coyote boldness categories across land cover, temporal and contextual variable categories. Reports are expressed as number (percentage); percentage is calculated as a function of the total number of reports within each category, so that each row sums to 100%.

| **Independent Variables** | | **Coyote Boldness Number of reports (percentage)** | | | |
|---|---|---|---|---|---|
| **Variable** | **Categories** | **Avoidance** | **Indifferent** | **Bold** | **Aggressive** |
| **Land Cover** | Natural | 143 (28.9%) | 178 (36%) | 110 (22.2%) | 64 (12.9%) |
| | Modified Open | 45 (20.3%) | 77 (34.7%) | 54 (24.3%) | 46 (20.7%) |
| | Mowed | 107 (22%) | 209 (42.9%) | 124 (25.5%) | 47 (9.7%) |
| | Residential | 606 (30.4%) | 888 (44.6%) | 341 (17.1%) | 158 (7.9%) |
| | Commercial | 92 (26.8%) | 180 (52.5%) | 53 (15.5%) | 18 (5.2%) |
| **Season** | Breeding | 352 (28.3%) | 611 (49.1%) | 189 (15.2%) | 93 (7.5%) |
| | Pup-Rearing | 185 (23.4%) | 275 (34.7%) | 193 (24.4%) | 139 (17.6%) |
| | Dispersal | 456 (30.3%) | 646 (43%) | 300 (20%) | 101 (6.7%) |
| **Time of Day** | Day | 490 (25.2%) | 889 (45.7%) | 365 (18.8%) | 201 (10.3%) |
| | Night | 449 (33%) | 544 (40%) | 264 (19.4%) | 103 (7.6%) |
| | Unknown | 54 (23%) | 99 (42.1%) | 53 (22.6%) | 29 (12.3%) |
| **Human Activity** | Cycling | 30 (52.6%) | 21 (36.8%) | 1 (1.8%) | 5 (8.8%) |
| | Driving | 174 (42.2%) | 228 (55.3%) | 3 (0.7%) | 7 (1.7%) |
| | Home/Yard | 221 (36.7%) | 257 (42.7%) | 52 (8.6%) | 72 (12%) |
| | Outdoor Activity | 33 (26.4%) | 49 (39.2%) | 29 (23.2%) | 14 (11.2%) |
| | Unknown | 246 (28.2%) | 441 (50.6%) | 99 (11.4%) | 86 (9.9%) |
| | Walking | 289 (19.6%) | 536 (36.4%) | 498 (33.8%) | 149 (10.1%) |
| **Vulnerable Individual** | Cat | 21 (17.5%) | 24 (20%) | 12 (10%) | 63 (52.5%) |
| | Child | 35 (23.3%) | 77 (51.3%) | 27 (18%) | 11 (7.3%) |
| | Dog | 274 (18.6%) | 494 (33.4%) | 495 (33.5%) | 214 (14.5%) |
| | Multiple | 35 (19.1%) | 77 (42.1%) | 45 (24.6%) | 26 (14.2%) |
| | Unknown | 628 (39%) | 860 (53.4%) | 103 (6.4%) | 19 (1.2%) |
| **Dog Leash Status** | Leashed | 33 (15.6%) | 65 (30.7%) | 83 (39.2%) | 31 (14.6%) |
| | Off-Leash | 38 (24.2%) | 33 (21%) | 35 (22.3%) | 51 (32.5%) |
| | Unknown | 199 (17.5%) | 410 (36.1%) | 399 (35.1%) | 128 (11.3%) |
| **Coyote Number** | One | 780 (31.6%) | 1086 (43.9%) | 425 (17.2%) | 181 (7.3%) |
| | Two | 124 (21.5%) | 240 (41.5%) | 138 (23.9%) | 76 (13.1%) |
| | Three | 35 (20.2%) | 66 (38.2%) | 51 (29.5%) | 21 (12.1%) |
| | More | 22 (16.4%) | 52 (38.8%) | 37 (27.6%) | 23 (17.2%) |
| | Unknown | 32 (17.5%) | 88 (48.1%) | 31 (16.9%) | 32 (17.5%) |
| **Health** | Healthy | 219 (35.7%) | 301 (49.1%) | 65 (10.6%) | 28 (4.6%) |
| | Unhealthy | 64 (35.4%) | 85 (47%) | 23 (12.7%) | 9 (5%) |
| | Unknown | 710 (25.9%) | 1146 (41.7%) | 594 (21.6%) | 296 (10.8%) |

**Table A2.2.** Distribution of human concern of coyote categories across land cover, temporal and contextual variable categories. Reports are expressed as number (percentage); percentage is calculated as a function of the total number of reports within each category, so that each row sums to 100%.

| Independent Variables | | Human Concern<br>Number of reports (category percentage) | | |
|---|---|---|---|---|
| **Variable** | **Categories** | **Negative** | **Neutral** | **Positive** |
| **Land Cover** | Natural | 80 (65%) | 25 (20.3%) | 18 (14.6%) |
| | Modified Open | 54 (68.4%) | 13 (16.5%) | 12 (15.2%) |
| | Mowed | 93 (66.9%) | 35 (25.2%) | 11 (7.9%) |
| | Residential | 425 (69.8%) | 99 (16.3%) | 85 (14%) |
| | Commercial | 66 (60%) | 23 (20.9%) | 21 (19.1%) |
| **Season** | Breeding | 260 (64%) | 85 (20.9%) | 61 (15%) |
| | Pup Rearing | 155 (67.4%) | 45 (19.6%) | 30 (13%) |
| | Dispersal | 303 (71.5%) | 65 (15.3%) | 56 (13.2%) |
| **Time of Day** | Day | 365 (65.6%) | 108 (19.4%) | 83 (14.9%) |
| | Night | 290 (68.1%) | 82 (19.2%) | 54 (12.7%) |
| | Unknown | 63 (80.8%) | 5 (6.4%) | 10 (12.8%) |
| **Human Activity** | Cycling | 2 (16.7%) | 8 (66.7%) | 2 (16.7%) |
| | Driving | 45 (46.4%) | 21 (21.6%) | 31 (32%) |
| | HomeYard | 214 (72.8%) | 42 (14.3%) | 38 (12.9%) |
| | OutdoorAct | 20 (76.9%) | 3 (11.5%) | 3 (11.5%) |
| | Unknown | 189 (69%) | 42 (15.3%) | 43 (15.7%) |
| | Walking | 248 (69.5%) | 79 (22.1%) | 30 (8.4%) |
| **Vulnerable Individual** | Cat | 32 (78%) | 2 (4.9%) | 7 (17.1%) |
| | Child | 91 (87.5%) | 10 (9.6%) | 3 (2.9%) |
| | Dog | 314 (77%) | 67 (16.4%) | 27 (6.6%) |
| | Multiple | 131 (92.3%) | 7 (4.9%) | 4 (2.8%) |
| | Unknown | 150 (41.1%) | 109 (29.9%) | 106 (29%) |
| **Dog Leash Status** | Leashed | 49 (75.4%) | 12 (18.5%) | 4 (6.2%) |
| | Off-Leash | 43 (87.8%) | 3 (6.1%) | 3 (6.1%) |
| | Unknown | 291 (79.7%) | 54 (14.8%) | 20 (5.5%) |
| **Coyote Number** | One | 396 (60.7%) | 141 (21.6%) | 115 (17.6%) |
| | Two | 138 (76.7%) | 26 (14.4%) | 16 (8.9%) |
| | Three | 62 (77.5%) | 9 (11.2%) | 9 (11.2%) |
| | More | 50 (80.6%) | 9 (14.5%) | 3 (4.8%) |
| | Unknown | 72 (83.7%) | 10 (11.6%) | 4 (4.7%) |
| **Health** | Healthy | 85 (38.3%) | 51 (23%) | 86 (38.7%) |
| | Unhealthy | 31 (77.5%) | 6 (15%) | 3 (7.5%) |
| | Unknown | 602 (75.4%) | 138 (17.3%) | 58 (7.3%) |

**Table A2.3.** Pearson's χ2 test of independence results examining if land cover, coyote season, time of day, or any contextual variables affected coyote boldness or human concern of coyotes.

| Variables Tested | *N*† | $\chi^2$ | df | *p* |
|---|---|---|---|---|
| Coyote boldness x Land cover | 3540 | 102.9 | 12 | 1.5E-16 |
| Coyote Boldness x Season | 3540 | 126.3 | 6 | 7.5E-25 |
| Coyote boldness x Time of day | 3305 | 30.1 | 3 | 1.3E-06 |
| Coyote boldness x Human activity | 3483 | 452.2 | 12 | 2.7E-92 |
| Coyote boldness x Vulnerable individual | 3540 | 916.8 | 12 | 1.4E-188 |
| Coyote boldness x Dog leash status | 1505 | 67.0 | 6 | 1.7E-12 |
| Coyote boldness x Number of coyotes | 3540 | 109.9 | 12 | 6.3E-18 |
| Coyote boldness x Coyote health | 3540 | 88.4 | 6 | 6.6E-17 |
| Human concern x Land cover | 1060 | 13.2 | 8 | 0.11 |
| Human concern x Season | 1060 | 6.1 | 4 | 0.2 |
| Human concern x Time of day | 982 | 1.1 | 2 | 0.58 |
| Human concern x Human activity | 1022 | 46.6 | 6 | 2.3E-08 |
| Human concern x Vulnerable individual | 1060 | 209.9 | 8 | 5.20E-41 |
| Human concern x Dog leash status | 452 | 3.67 | 2 | 0.16 |
| Human concern x Number of coyotes | 1060 | 42.0 | 8 | 1.3E-06 |
| Human concern x Coyote health | 1060 | 164.5 | 4 | 1.6E-34 |
| Human concern x Coyote boldness | 653 | 56.3 | 6 | 2.5E-10 |

† *N* is the number of reports available for each test

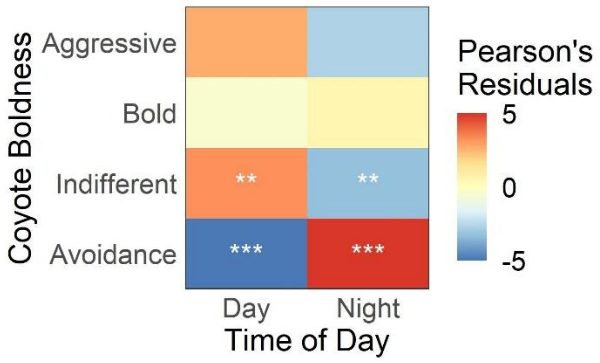


**Figure A2.1.** Relationship between coyote boldness and time of day. Colors represent Pearson's residual values calculated post-hoc from chi square tests, with positive values (red) indicating positive relationships and negative values (blue) indicating negative relationships. Significance is indicated by asterisks (* p < 0.05, ** p <0.01, *** p < 0.001).
