## Appendix 3 for "A ten-year community reporting database reveals rising coyote boldness and associated human concern in Edmonton, Canada"

### APPENDIX 3. ORDERED LOGISTIC REGRESSION MODELLING

**Table A3.1.** Pearson's chi-square test of independence outputs testing the relationship between season, time of day, and contextual variables.

| **Variables Tested** | ***N*†** | $\chi^2$ | **df** | ***p*** |
|---|---|---|---|---|
| Season x Human activity | 9134 | 66.5 | 10 | 2.1E-10 |
| Season x Vulnerable individual | 9134 | 77.0 | 8 | 1.9E-13 |
| Season x Dog leash status | 2202 | 23.6 | 6 | 6.2E-04 |
| Season x Number of coyotes | 9134 | 95.5 | 8 | 3.5E-17 |
| Season x Coyote health | 9134 | 19.5 | 4 | 6.4E-04 |
| Season x Time of day | 8474 | 298.9 | 2 | 1.3E-65 |
| Time of day x Human activity | 8474 | 92.7 | 5 | 1.8E-18 |
| Time of day x Vulnerable individual | 8474 | 39.2 | 4 | 6.3E-08 |
| Time of day x Dog leash status | 2029 | 51.1 | 3 | 4.6E-11 |
| Time of day x Number of coyotes | 8474 | 240.8 | 4 | 6.3E-51 |
| Time of day x Coyote health | 8474 | 155.3 | 2 | 1.9E-34 |
| Human activity x Vulnerable individual | 1262 | 89.8 | 6 | 3.4E-17 |
| Human activity x Dog leash status | 1882 | 60.8 | 4 | 2.0E-12 |
| Human activity x Number of coyotes | 9059 | 624.4 | 16 | 1.5E-122 |
| Human activity x Coyote health | 9059 | 108.1 | 8 | 9.2E-20 |
| Vulnerable individual x Dog leash status | 1958 | 13.6 | 2 | 1.1E-03 |
| Vulnerable individual x Number of coyotes | 9134 | 193.5 | 16 | 1.6E-32 |
| Vulnerable individual x Coyote health | 9134 | 40.9 | 8 | 2.2E-06 |
| Dog leash status x Number of coyotes | 9134 | 193.5 | 16 | 1.6E-32 |
| Dog leash status x Coyote health | 2202 | 5.4 | 6 | 5.0E-01 |
| Number of coyotes x Coyote health | 9134 | 464.8 | 8 | 2.5E-95 |

† *N* is the number of reports available for each test

**Table A3.2** The outputs from univariate ordinal regression models aiming to determine the best-fit scale for measuring land cover variables and building density, showing AIC values and the difference between the AIC value of each univariate model and the null model.

| **Coyote Boldness** | | | **Human Concern** | | |
|---|---|---|---|---|---|
| Variable (scale)† | AIC | $\Delta AIC_{null}$ | Variable (scale)† | AIC | $\Delta AIC_{null}$ |
| Building Density (100m) | 8865.2 | -52.1 | Building Density (100m) | 1805.2 | 0.7 |
| **Building Density (200m)** | **8860.7** | **-56.6** | **Building Density (200m)** | **1804.5** | **0.0** |
| Building Density (400m) | 8868.1 | -49.2 | Building Density (400m) | 1804.8 | 0.3 |
| Building Density (800m) | 8885.2 | -32.1 | Building Density (800m) | 1806.3 | 1.8 |
| Building Density (1600m) | 8899.3 | -18.1 | Building Density (1600m) | 1805.6 | 1.0 |
| **Natural (100m)** | **8911.2** | **-6.1** | Natural (100m) | 1803.5 | -1.0 |
| Natural (200m) | 8912.5 | -4.8 | Natural (200m) | 1803.7 | -0.8 |
| Natural (400m) | 8911.9 | -5.4 | Natural (400m) | 1804.9 | 0.4 |
| Natural (800m) | 8911.4 | -6.0 | Natural (800m) | 1804.6 | 0.1 |
| Natural (1600m) | 8918.0 | 0.6 | **Natural (1600m)** | **1802.2** | **-2.3** |
| Modified Open (100m) | 8892.7 | -24.6 | Modified Open (100m) | 1806.5 | 2.0 |
| Modified Open (200m) | 8889.9 | -27.4 | Modified Open (200m) | 1806.5 | 2.0 |
| **Modified Open (400m)** | **8889.1** | **-28.2** | Modified Open (400m) | 1806.5 | 2.0 |
| Modified Open (800m) | 8898.2 | -19.1 | Modified Open (800m) | 1806.5 | 2.0 |
| Modified Open (1600m) | 8899.9 | -17.4 | **Modified Open (1600m)** | **1802.1** | **-2.4** |
| **Mowed (100m)** | **8906.5** | **-10.8** | Mowed (100m) | 1805.6 | 1.1 |
| Mowed (200m) | 8914.2 | -3.1 | Mowed (200m) | 1806.5 | 2.0 |
| Mowed (400m) | 8919.3 | 1.9 | Mowed (400m) | 1805.1 | 0.6 |
| Mowed (800m) | 8918.0 | 0.7 | **Mowed (800m)** | **1801.1** | **-3.4** |
| Mowed (1600m) | 8918.6 | 1.3 | Mowed (1600m) | 1805.3 | 0.8 |
| Commercial (100m) | 8916.0 | -1.3 | **Commercial (100m)** | **1802.3** | **-2.2** |
| **Commercial (200m)** | **8915.7** | **-1.6** | Commercial (200m) | 1804.3 | -0.2 |
| Commercial (400m) | 8915.8 | -1.5 | Commercial (400m) | 1804.9 | 0.4 |
| Commercial (800m) | 8917.0 | -0.3 | Commercial (800m) | 1805.5 | 1.0 |
| Commercial (1600m) | 8918.1 | 0.8 | Commercial (1600m) | 1804.4 | -0.1 |
| **Residential (100m)** | **8877.4** | **-40.0** | Residential (100m) | 1803.7 | -0.8 |
| Residential (200m) | 8890.8 | -26.6 | Residential (200m) | 1801.4 | -3.1 |
| Residential (400m) | 8903.4 | -13.9 | Residential (400m) | 1798.7 | -5.8 |
| Residential (800m) | 8910.3 | -7.1 | **Residential (800m)** | **1798.3** | **-6.2** |
| Residential (1600m) | 8912.9 | -4.4 | Residential (1600m) | 1801.6 | -2.9 |

† Bolded values indicate the best-fit scale (lowest AIC value)

**Table A3.3.** Spearman's correlation coefficients between variables used for ordinal regression models of coyote boldness towards humans. For variable pairs with r > 0.6, only the variable with the lowest AIC in univariate models was retained for further analysis.

| | Road Distance Decay | Modified Open (400m) | Natural (100m) | Mowed (100m) | Building Density (200m) | Commercial (200m) | Residential (100m) |
|---|---|---|---|---|---|---|---|
| Road Distance Decay | - | -0.12 | -0.18 | -0.13 | 0.50 | 0.14 | 0.58 |
| Modified Open (400m) | - | - | 0.02 | -0.34 | -0.14 | -0.11 | -0.06 |
| Natural (100m) | - | - | - | -0.28 | -0.39 | -0.31 | -0.23 |
| Mowed (100m) | - | - | - | - | 0.00 | 0.02 | -0.15 |
| Building Density (200m) | - | - | - | - | - | 0.16 | **0.61** |
| Commercial (200m) | - | - | - | - | - | - | -0.17 |
| Residential (100m) | - | - | - | - | - | - | - |

**Table A3.4.** Spearman's correlation coefficients between variables used for ordinal regression models of human concern of coyotes. For variable pairs with r > 0.6 (bold), only the variable with the lower AIC in univariate models was retained for further analysis.

| | Modified Open (1600m) | Mowed (800m) | Natural (1600m) | Commercial (100m) | Residential (800m) |
|---|---|---|---|---|---|
| Modified Open (1600m) | - | -0.42 | 0.062 | -0.15 | 0.024 |
| Mowed (800m) | - | - | -0.03 | -0.017 | -0.02 |
| Natural (1600m) | - | - | - | -0.19 | -0.32 |
| Commercial (100m) | - | - | - | - | -0.13 |
| Residential (800m) | - | - | - | - | - |

**Table A3.5.** The variables included in global models examining the factors affecting coyote boldness towards humans and human concern of coyotes

| Global Model | Variables | df | AIC | ΔAIC$_{null}$ |
|---|---|---|---|---|
| **Coyote Boldness** | RoadDistDecay + BuildingDensity200m + Natural100m + ModifiedOpen400m + Mowed100m + Commercial200m + Season + Natural100m*Season + ModifiedOpen*Season + Year + RoadDistDecay*Year + BuildingDensity200m *Year + Natural100m*Year + ModifiedOpen400m*Year + Mowed100m*Year + Commercial200m*Year | 22 | 8691.3 | -226.1 |
| **Human Concern** | Natural1600m + Modified_Open1600m + Mowed800m + Commercial100m + Residential800m + Season + Year + Year*Natural1600m + Year*Modified_Open1600m + Year*Mowed800m + Year*Commercial100m + Year*Residential800m | 15 | 1797.2 | 8.2 |

**Table A3.6.** The coefficients, confidence intervals and model parameters from the top-ranked ordinal regression models assessing coyote boldness towards humans.

| Variables† ß (2.5% C.I. , 97.5% C.I.) | | | | | | | | | | | | | | Model Parameters | | |
|---|---|---|---|---|---|---|---|---|---|---|---|---|---|---|---|---|
| **BUILD** | **MOD** | **MOW** | **ROAD** | **SEAS(D)** | **SEAS(P)** | **YEAR** | **MOD: SEAS(D)** | **MOD: SEAS(P)** | **MOW: YEAR** | **MOD: YEAR** | **ROAD: YEAR** | **BUILD: YEAR** | **COM: YEAR** | $AIC_c$ | $\Delta AIC_c$ | wgt$AIC_c$ |
| -0.13 (-0.2, -0.05) | -0.02 (-0.13, 0.09) | 0.09 (0.02, 0.16) | -0.11 (-0.18, -0.03) | 0.03 (-0.1 , 0.17) | 0.03 (-0.1 , 0.17) | 0.29 (0.22 , 0.35) | 0.12 (-0.02 , 0.27) | 0.36 (0.19 , 0.52) | NA | NA | NA | NA | NA | 8677.6 | 0 | 0.14 |
| -0.13 (-0.2, -0.05) | -0.02 (-0.14, 0.09) | 0.09 (0.02, 0.15) | -0.11 (-0.18, -0.03) | 0.03 (-0.11 , 0.17) | 0.03 (-0.11 , 0.17) | 0.28 (0.22 , 0.35) | 0.13 (-0.02 , 0.27) | 0.36 (0.2 , 0.52) | 0.04 (-0.02 , 0.1) | NA | NA | NA | NA | 8677.7 | 0.1 | 0.14 |
| -0.13 (-0.2, -0.05) | -0.02 (-0.13, 0.09) | 0.09 (0.02, 0.15) | -0.11 (-0.18 , -0.03) | 0.03 (-0.1 , 0.17) | 0.03 (-0.1 , 0.17) | 0.29 (0.22 , 0.35) | 0.12 (-0.02 , 0.27) | 0.36 (0.2 , 0.52) | NA | -0.04 (-0.1 , 0.02) | NA | NA | NA | 8677.7 | 0.13 | 0.13 |
| -0.13 (-0.2, -0.05) | -0.02 (-0.14, 0.09) | 0.09 (0.02, 0.15) | -0.11 (-0.18 , -0.03) | 0.03 (-0.11 , 0.17) | 0.03 (-0.11 , 0.17) | 0.29 (0.22 , 0.35) | 0.13 (-0.02 , 0.27) | 0.36 (0.2 , 0.53) | 0.04 (-0.03 , 0.1) | -0.03 (-0.09 , 0.03) | NA | NA | NA | 8678.5 | 0.92 | 0.09 |
| -0.13 (-0.2, -0.05) | -0.02 (-0.13, 0.09) | 0.08 (0.02, 0.15) | -0.11 (-0.18 , -0.04) | 0.03 (-0.11 , 0.17) | 0.03 (-0.11 , 0.17) | 0.28 (0.22 , 0.35) | 0.12 (-0.02 , 0.27) | 0.36 (0.2 , 0.52) | 0.05 (-0.01 , 0.11) | NA | 0.04 (-0.03 , 0.1) | NA | NA | 8678.5 | 0.93 | 0.09 |
| -0.12 (-0.2, -0.05) | -0.02 (-0.13, 0.09) | 0.09 (0.02, 0.15) | -0.11 (-0.18 , -0.03) | 0.04 (-0.1 , 0.17) | 0.04 (-0.1 , 0.17) | 0.28 (0.22 , 0.35) | 0.12 (-0.02 , 0.27) | 0.35 (0.19 , 0.52) | NA | NA | 0.02 (-0.04 , 0.08) | NA | NA | 8679.1 | 1.51 | 0.07 |
| -0.13 (-0.2, -0.05) | -0.03 (-0.14, 0.09) | 0.09 (0.02, 0.15) | -0.11 (-0.18 , -0.03) | 0.03 (-0.11 , 0.17) | 0.03 (-0.11 , 0.17) | 0.28 (0.22 , 0.35) | 0.13 (-0.02 , 0.27) | 0.36 (0.2 , 0.52) | 0.05 (-0.01 , 0.11) | NA | NA | 0.02 (-0.04 , 0.09) | NA | 8679.1 | 1.54 | 0.07 |
| -0.13 (-0.2, -0.05) | -0.02 (-0.13, 0.09) | 0.09 (0.02, 0.15) | -0.11 (-0.18 , -0.03) | 0.04 (-0.1 , 0.17) | 0.04 (-0.1 , 0.17) | 0.29 (0.22 , 0.35) | 0.12 (-0.02 , 0.27) | 0.36 (0.19 , 0.52) | NA | NA | NA | 0.01 (-0.05 , 0.08) | NA | 8679.4 | 1.78 | 0.06 |

| | | | | | | | | | | | | | | | | |
|---|---|---|---|---|---|---|---|---|---|---|---|---|---|---|---|---|
| -0.12 (-0.2, -0.05) | -0.02 (-0.13, 0.09) | 0.09 (0.02, 0.15) | -0.11 (-0.18 , -0.03) | 0.03 (-0.1 , 0.17) | 0.03 (-0.1 , 0.17) | 0.28 (0.22 , 0.35) | 0.12 (-0.02 , 0.27) | 0.36 (0.19 , 0.52) | NA | NA | NA | NA | -0.01 (-0.07 , 0.05) | 8679.4 | 1.84 | 0.06 |
| -0.12 (-0.2, -0.05) | -0.02 (-0.13, 0.09) | 0.09 (0.02, 0.15) | -0.11 (-0.18 , -0.03) | 0.03 (-0.11 , 0.17) | 0.03 (-0.11 , 0.17) | 0.29 (0.22 , 0.35) | 0.12 (-0.02 , 0.27) | 0.36 (0.2 , 0.52) | NA | -0.04 (-0.1 , 0.02) | NA | NA | -0.01 (-0.07 , 0.05) | 8679.5 | 1.95 | 0.05 |
| -0.12 (-0.2, -0.05) | -0.02 (-0.14, 0.09) | 0.09 (0.02, 0.15) | -0.11 (-0.18 , -0.03) | 0.03 (-0.11 , 0.17) | 0.03 (-0.11 , 0.17) | 0.28 (0.22 , 0.35) | 0.13 (-0.02 , 0.27) | 0.36 (0.2 , 0.52) | 0.04 (-0.02 , 0.1) | NA | NA | NA | -0.01 (-0.07 , 0.05) | 8679.5 | 1.96 | 0.05 |
| -0.13 (-0.2, -0.05) | -0.02 (-0.13, 0.09) | 0.09 (0.02, 0.15) | -0.11 (-0.18, -0.03) | 0.03 (-0.1 , 0.17) | 0.03 (-0.1 , 0.17) | 0.29 (0.22 , 0.35) | 0.12 (-0.02 , 0.27) | 0.36 (0.2 , 0.52) | NA | -0.04 (-0.1 , 0.02) | 0.01 (-0.05 , 0.08) | NA | NA | 8679.5 | 1.99 | 0.05 |

† BUILD = Building Density (200m), MOD = Modified Open (400m), MOW = Mowed (100m), ROAD = Road Distance Decay, SEAS(D) = Season (Dispersal), SEAS(P) = Season (Pup rearing), YEAR = Year, COM = Commercial (200m)

**Table A3.7.** The coefficients, confidence intervals and model parameters from the top-ranked ordinal regression models assessing human concern of coyotes.

| Variables† ß (2.5% C.I. , 97.5 C.I.) | | | | | | | | | | Model Parameters | | |
|---|---|---|---|---|---|---|---|---|---|---|---|---|
| **RES** | **YEAR** | **RES: YEAR** | **MOD** | **SEAS(D)** | **SEAS(P)** | **MOD: YEAR** | **NAT** | **MOW** | **COM** | **AIC$_c$** | **ΔAIC$_c$** | **wgtAIC$_c$** |
| 0.17 (0.04 , 0.3) | 0.14 (0.01 , 0.27) | -0.12 (-0.25 , 0) | 0.16 (0.03 , 0.3) | 0.33 (0.04 , 0.62) | 0.18 (-0.16 , 0.52) | NA | NA | NA | NA | 1788.9 | 0.00 | 0.10 |
| 0.17 (0.04 , 0.29) | 0.13 (0 , 0.26) | -0.13 (-0.25 , 0) | 0.16 (0.03 , 0.3) | 0.32 (0.03 , 0.61) | 0.17 (-0.17 , 0.51) | -0.09 (-0.23 , 0.04) | NA | NA | NA | 1789.1 | 0.30 | 0.08 |
| 0.17 (0.04 , 0.29) | 0.13 (0 , 0.26) | -0.13 (-0.26 , -0.01) | 0.15 (0.02 , 0.29) | NA | NA | -0.1 (-0.24 , 0.03) | NA | NA | NA | 1789.7 | 0.84 | 0.06 |
| 0.14 (0 , 0.28) | 0.13 (0 , 0.26) | -0.13 (-0.25 , 0) | 0.16 (0.03 , 0.3) | 0.33 (0.04 , 0.62) | 0.18 (-0.16 , 0.52) | NA | -0.07 (-0.21 , 0.06) | NA | NA | 1789.8 | 0.94 | 0.06 |
| 0.17 (0.04 , 0.3) | 0.14 (0.01 , 0.26) | -0.13 (-0.25 , 0) | 0.15 (0.02 , 0.28) | NA | NA | NA | NA | NA | NA | 1789.9 | 0.95 | 0.06 |
| 0.16 (0.03 , 0.29) | 0.14 (0.01 , 0.26) | -0.12 (-0.25 , 0.01) | 0.13 (-0.02 , 0.28) | 0.33 (0.04 , 0.62) | 0.18 (-0.16 , 0.52) | NA | NA | -0.07 (-0.21 , 0.07) | NA | 1789.9 | 1.05 | 0.06 |
| 0.14 (0 , 0.28) | 0.13 (0 , 0.26) | -0.13 (-0.26 , -0.01) | 0.16 (0.03 , 0.3) | 0.32 (0.03 , 0.61) | 0.17 (-0.17 , 0.51) | -0.09 (-0.22 , 0.05) | -0.07 (-0.2 , 0.07) | NA | NA | 1790.2 | 1.38 | 0.05 |
| 0.15 (0.01 , 0.29) | 0.14 (0.01 , 0.27) | -0.13 (-0.25 , 0) | 0.15 (0.01 , 0.29) | 0.33 (0.04 , 0.62) | 0.17 (-0.16 , 0.51) | NA | NA | NA | -0.05 (-0.19 , 0.08) | 1790.3 | 1.45 | 0.05 |
| 0.16 (0.03 , 0.29) | 0.13 (0 , 0.26) | -0.12 (-0.25 , 0) | 0.13 (-0.01 , 0.28) | 0.32 (0.03 , 0.61) | 0.17 (-0.16 , 0.51) | -0.09 (-0.22 , 0.05) | NA | -0.07 (-0.2 , 0.07) | NA | 1790.3 | 1.48 | 0.05 |
| 0.09 (-0.06 , 0.25) | 0.13 (0.01 , 0.26) | -0.13 (-0.26 , -0.01) | 0.15 (0.01 , 0.29) | 0.33 (0.04 , 0.62) | 0.17 (-0.16 , 0.52) | NA | -0.1 (-0.25 , 0.04) | NA | -0.09 (-0.23 , 0.06) | 1790.4 | 1.57 | 0.04 |

| | | | | | | | | | | | | |
|---|---|---|---|---|---|---|---|---|---|---|---|---|
| 0.14 (0.01 , 0.28) | 0.13 (0.01 , 0.26) | -0.13 (-0.26 , -0.01) | 0.15 (0.02 , 0.29) | 0.31 (0.02 , 0.61) | 0.17 (-0.17 , 0.51) | -0.09 (-0.23 , 0.04) | NA | NA | -0.06 (-0.19 , 0.08) | 1790.5 | 1.67 | 0.04 |
| 0.18 (0.06 , 0.31) | 0.15 (0.02 , 0.28) | NA | 0.16 (0.03 , 0.3) | 0.34 (0.05 , 0.63) | 0.19 (-0.15 , 0.53) | NA | NA | NA | NA | 1790.6 | 1.69 | 0.04 |
| 0.06 (-0.1 , 0.21) | 0.13 (0 , 0.26) | -0.13 (-0.26 , 0) | NA | 0.31 (0.02 , 0.6) | 0.17 (-0.16 , 0.52) | NA | -0.12 (-0.26 , 0.03) | -0.13 (-0.26 , -0.01) | -0.12 (-0.26 , 0.02) | 1790.6 | 1.84 | 0.04 |
| 0.13 (-0.01 , 0.27) | 0.13 (0 , 0.26) | -0.12 (-0.25 , 0) | 0.13 (-0.02 , 0.28) | 0.33 (0.04 , 0.62) | 0.18 (-0.16 , 0.52) | NA | -0.08 (-0.21 , 0.06) | -0.07 (-0.21 , 0.07) | NA | 1790.7 | 1.88 | 0.04 |
| 0.15 (0.02 , 0.28) | 0.14 (0.01 , 0.26) | -0.12 (-0.25 , 0.01) | NA | 0.31 (0.02 , 0.6) | 0.18 (-0.16 , 0.52) | NA | NA | -0.12 (-0.25 , 0) | NA | 1790.8 | 1.91 | 0.04 |
| 0.16 (0.03 , 0.29) | 0.13 (0.01 , 0.26) | -0.12 (-0.25 , 0) | 0.12 (-0.03 , 0.27) | NA | NA | NA | NA | -0.07 (-0.21 , 0.07) | NA | 1790.8 | 1.94 | 0.04 |
| 0.09 (-0.06 , 0.25) | 0.13 (0 , 0.26) | -0.14 (-0.26 , -0.01) | 0.15 (0.01 , 0.29) | 0.32 (0.03 , 0.61) | 0.17 (-0.17 , 0.51) | -0.09 (-0.23 , 0.05) | -0.1 (-0.24 , 0.05) | NA | -0.09 (-0.23 , 0.06) | 1790.7 | 1.95 | 0.04 |
| 0.14 (0.01 , 0.28) | 0.13 (0 , 0.26) | -0.13 (-0.26 , 0) | 0.15 (0.02 , 0.29) | NA | NA | NA | -0.07 (-0.2 , 0.07) | NA | NA | 1790.9 | 1.97 | 0.04 |
| 0.16 (0.03 , 0.29) | 0.13 (0 , 0.26) | -0.13 (-0.26 , 0) | 0.12 (-0.03 , 0.27) | NA | NA | -0.1 (-0.23 , 0.04) | NA | -0.07 (-0.21 , 0.07) | NA | 1790.8 | 1.98 | 0.04 |
| 0.14 (0.01 , 0.28) | 0.13 (0 , 0.25) | -0.13 (-0.26 , -0.01) | 0.15 (0.02 , 0.29) | NA | NA | -0.1 (-0.23 , 0.04) | -0.06 (-0.2 , 0.07) | NA | NA | 1790.9 | 2.00 | 0.04 |

† MOD = Modified Open (1600m), RES = Residential (800m), SEAS(D) = Season (dispersal), SEAS(P) = Season (pup rearing), YEAR = Year, NAT = Natural (1600m), MOW = Mowed (800m), COM = Commercial

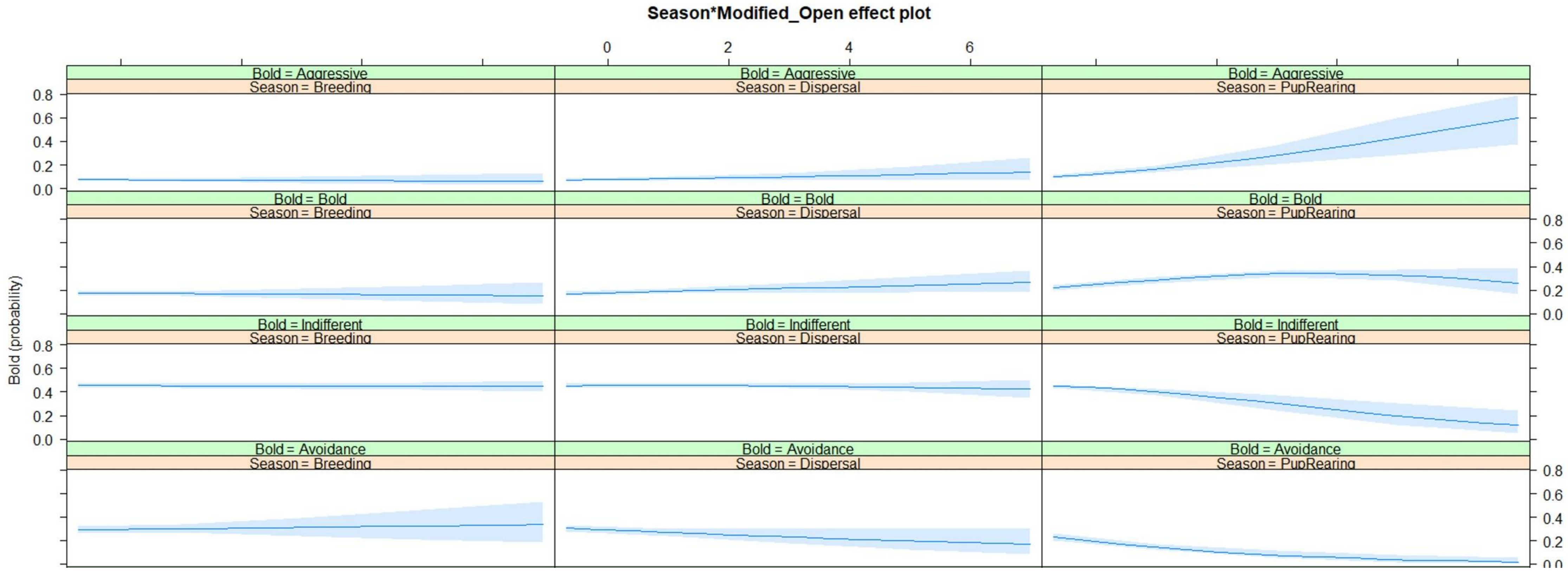


**Figure A3.1.** Interaction plot between season and modified open land cover (within 400m) for models of coyote boldness. The plot was generated using the *Effect* function from the package effects (Fox and Hong 2009) on an ordered logistic regression model created using the function *polr* from the package MASS (Venables and Ripley 2002).

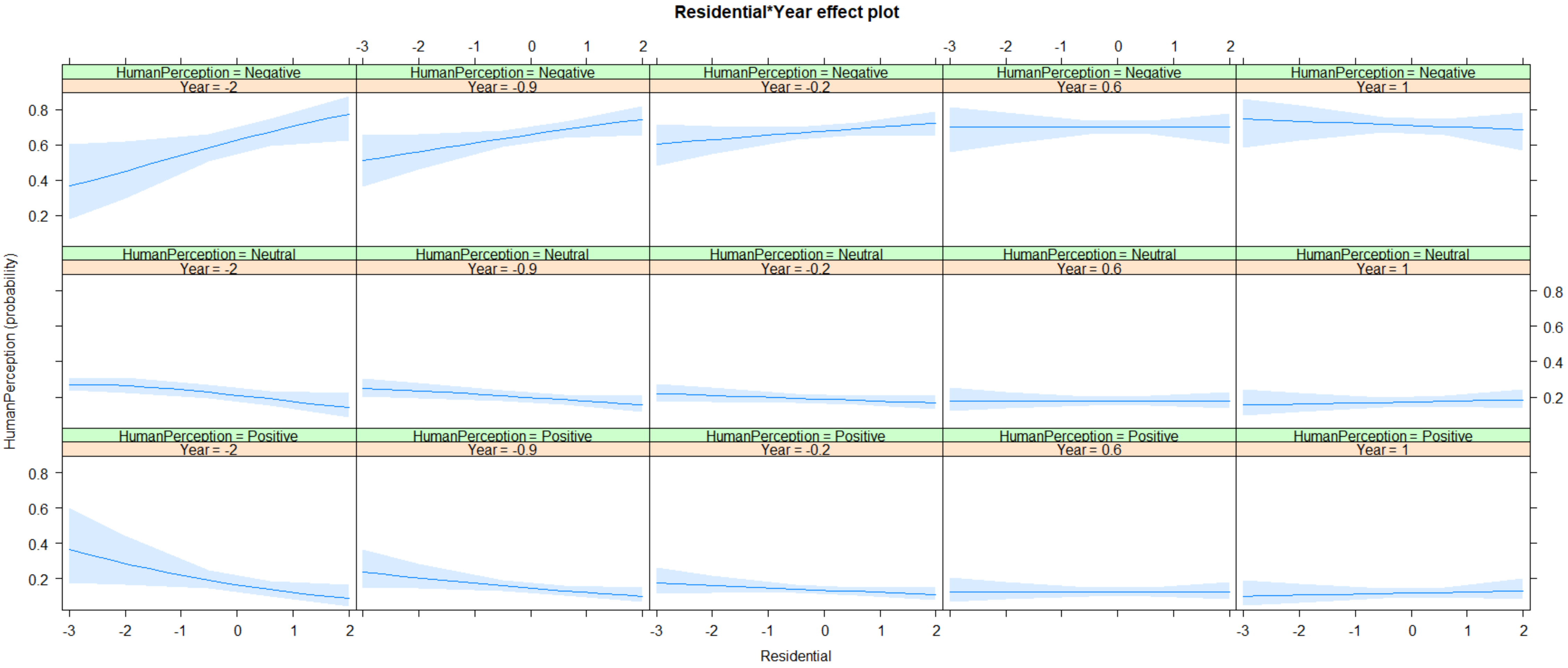


**Figure A3.2.** Interaction plot between year and the proportion of residential area within 800 m for human concern models. The plot was generated using the *Effect* function from the package effects (Fox and Hong 2009) on an ordered logistic regression model created using the function *polr* from the package MASS (Venables and Ripley 2002).
